# Variant Curation Expert Panel Recommendations for RYR1 Pathogenicity Assertions in Malignant Hyperthermia Susceptibility

**DOI:** 10.1101/2020.11.29.402768

**Authors:** Jennifer J. Johnston, Robert T. Dirksen, Thierry Girard, Stephen G. Gonsalves, Phil M. Hopkins, Sheila Riazi, Louis A. Saddic, Nyamkhishig Sambuughin, Richa Saxena, Kathryn Stowell, James Weber, Henry Rosenberg, Leslie G. Biesecker

## Abstract

**Purpose:** Prevention of malignant hyperthermia (MH) requires an understanding of *RYR1* variant pathogenicity to assess the risk of exposure to triggering agents. Personalized medicine, especially secondary findings and eventually genomic screening, will contribute toward this goal.

**Methods:** We specified ACMG/AMP criteria for variant interpretation for *RYR1* and MH. Proposed rules were piloted on 84 variants. We applied quantitative evidence calibration for several criteria using likelihood ratios based on the Bayesian framework.

**Results:** Seven ACMG/AMP criteria were adopted without changes, ten were adopted with *RYR1*-specific modifications, and nine were dropped. The *in silico* (PP3 and BP4) and hot spot criteria (PM1) were evaluated quantitatively. REVEL gave an OR of 23:1 for PP3 and 16:1 for BP4 using trichotomized cut-offs of >0.85 (pathogenic) and <0.5 (benign). The PM1 hotspot criterion had an OR of 24:1. PP3 and PM1 were implemented at moderate strength. Applying the revised ACMG criteria to 44 recognized MH variants, 30 were assessed as pathogenic, 12 as likely pathogenic, and two as VUS.

**Conclusion:** Curation of these variants will facilitate interpretation of *RYR1*/MH genomic testing results, which is especially important for secondary findings analyses. Our approach to quantitatively calibrating criteria are generalizable to other variant curation expert panels.

## INTRODUCTION

Malignant hyperthermia susceptibility (MHS) is a potentially lethal inherited disorder of skeletal muscle calcium signaling, predisposing individuals to a hypermetabolic reaction triggered by exposure to inhalational anesthetics or depolarizing muscle relaxants such as succinylcholine.^1,2^ Inheritance of MHS is predominantly autosomal dominant, although autosomal recessive inheritance has been reported^3^ and non-Mendelian models proposed.^4^ Variants in *RYR1* (MIM:180901; MHS1, MIM:145600) and *CACNA1S* (MIM:114208; MHS5, MIM:601887) have been identified as associated with MH, and both genes are in the American College of Medical Genetics and Genomics (ACMG) return of secondary findings recommendations.^5,6^ *RYR1* variants account for ~76% of MH events while ~1%^7^ are attributable to *CACNA1S* and <1% are attributable to *STAC3* (MIM:615521; Bailey-Bloch myopathy, MIM:255995). Four additional loci have been mapped (MHS2, MIM:154275; MHS3, MIM:154276; MHS4, MIM:600467; MHS6, MIM:601888). *RYR1* has a complex gene-to-phenotype relationship, being associated with several apparently distinct disorders and both autosomal dominant and autosomal recessive inheritance. Overlapping conditions include central core disease (CCD, MIM:117000) and King-Denborough syndrome (MIM:145600) and individuals with these disorders may be at risk for MH. Generally, these disorders result from monoallelic *RYR1* variants while biallelic variants cause other myopathies including neuromuscular disease with uniform type 1 fiber (MIM:117000) and minicore myopathy with external ophthalmoplegia (MIM:255320), however, this correlation is evolving.^8^

Interpretation of *RYR1* variants is complicated by variable expressivity, reduced penetrance and high alleleic heterogeneity. While the European Malignant Hyperthermia Group (EMHG; http://www.emhg.org/home/) has assessed 48 *RYR1* variants as diagnostic of MHS, over 165 additional variants have been reported as disease mutations/pathogenic/likely pathogenic for MH in the literature and databases including HGMD^9,10^ and ClinVar.^13^ While the ACMG/AMP guidelines provided general criteria that can be used to assess variants, many of the criteria require adaptation to be accurately applied. As part of ClinGen, we convened an *RYR1-*related Malignant Hyperthermia variant curation expert panel (https://clinicalgenome.org/affiliation/50038/) to adapt the general ACMG/AMP pathogenicity guidelines to autosomal dominantly inherited *RYR1/*MH, with gene-specific recommendations, to improve interpretation of *RYR1* variants.

We first reviewed each ACMG/AMP criterion to determine their applicability to autosomal dominantly inherited *RYR1/*MH and then adapted them with gene/disease specific guidelines, if appropriate. We piloted these guidelines on 84 variants – 44 variants from the EMHG list of diagnostic variants and 40 variants with MH pathogenicity assertions in ClinVar.

## METHODS

### ClinGen’s RYR1/MH Expert Panel

The *RYR1/*MH expert panel (EP) is composed of clinical molecular geneticists, clinical geneticists, anesthesiologists, biochemists, and physiologists to provide a balance of expertise relevant to *RYR1* variant interpretation. The *RYR1*/MH EP met monthly via conference calls over a two-year period.

### Evaluation and Adaptation of the ACMG Pathogenicity Guidelines

The general ACMG/AMP pathogenicity guidelines were evaluated for relevance to autosomal dominantly inherited *RYR1/*MH and criteria that were not relevant were dropped. ClinGen-recommended amendments to the ACMG/AMP criteria were incorporated when applicable. Lastly, applicable criteria were further assessed to determine if gene-specific recommendations were warranted. Proposed changes were discussed amongst the full EP by means of emails and conference calls to develop consensus. Draft rules were piloted on a subset of *RYR1* variants representing the EMHG diagnostic variant list. Individual panel members scored variants using the draft guidelines and variant interpretations were presented to the full panel. Areas of disagreement were used to refine the draft guidelines.

### Data Collection Methods

Population data for each variant were ascertained from gnomAD.^11^ REVEL scores were used for bioinformatic predictions for single nucleotide variants (SNVs).^12^ The literature was searched for data relevant to each variant including case information and functional data. For case information, the number of unrelated probands with either a personal or family history of an MH event was recorded (see supplemental information). Care was taken to avoid double counting cases reported multiple times. Reports were examined for instances of *de novo* inheritance and/or segregation.

### Pathogenicity Assessment

Revised ACMG/AMP criteria were used to assess 44 EMHG MH diagnostic variants. Four of 48 EMHG variants were excluded because they were only associated with *RYR1*-related myopathies and not MH. An additional 40 ClinVar *RYR1* variants were also assessed. Individual criteria were weighted based on available evidence and then weighted criteria were combined using the Bayesian framework for variant scoring.^13^

## RESULTS AND DISCUSSION

The ACMG/AMP guidelines are generic and broadly useful for all genes and disorders. These generic rules may over- or under-estimate evidence for any specific gene and must be adapted for specific implementations. As an EP, we suggest guidelines to be used/dropped, guidelines to be refined, and weight adjustments where appropriate. Summary of revised guidelines are in Table 1 and a full description of the revised guidelines is in Table S1 with gene/disease specific adaptations highlighted below.

**Table 1.**
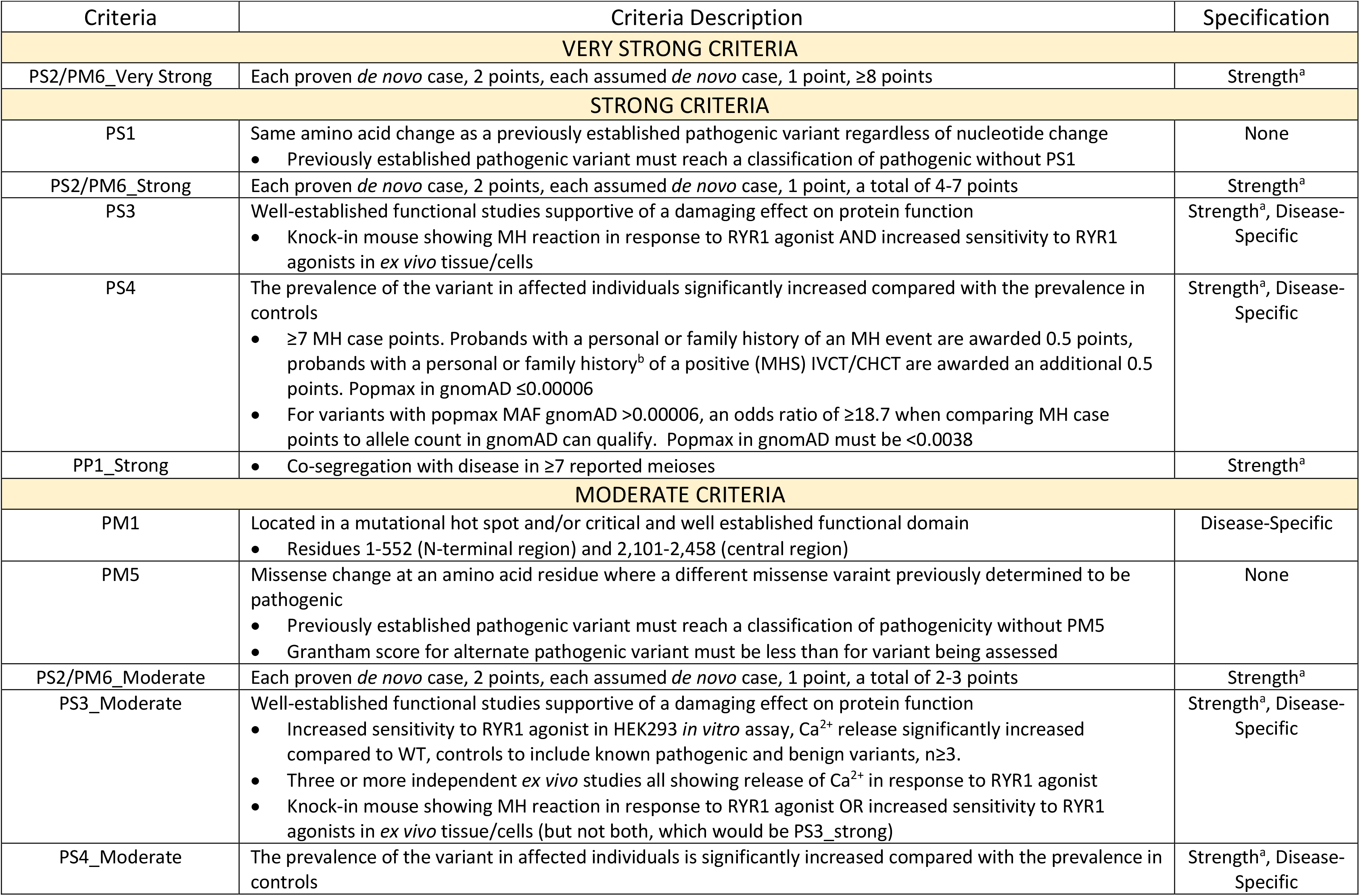
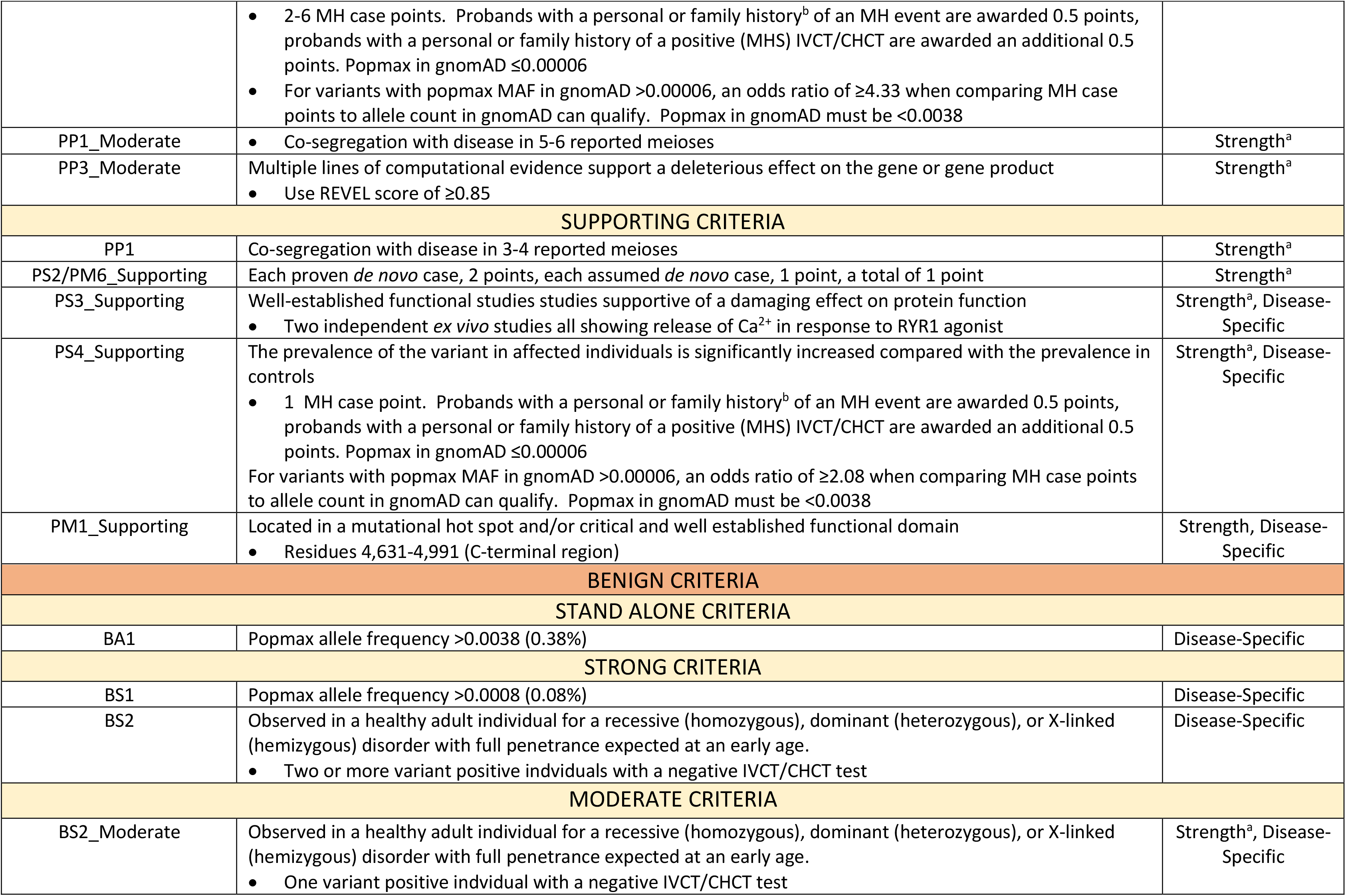
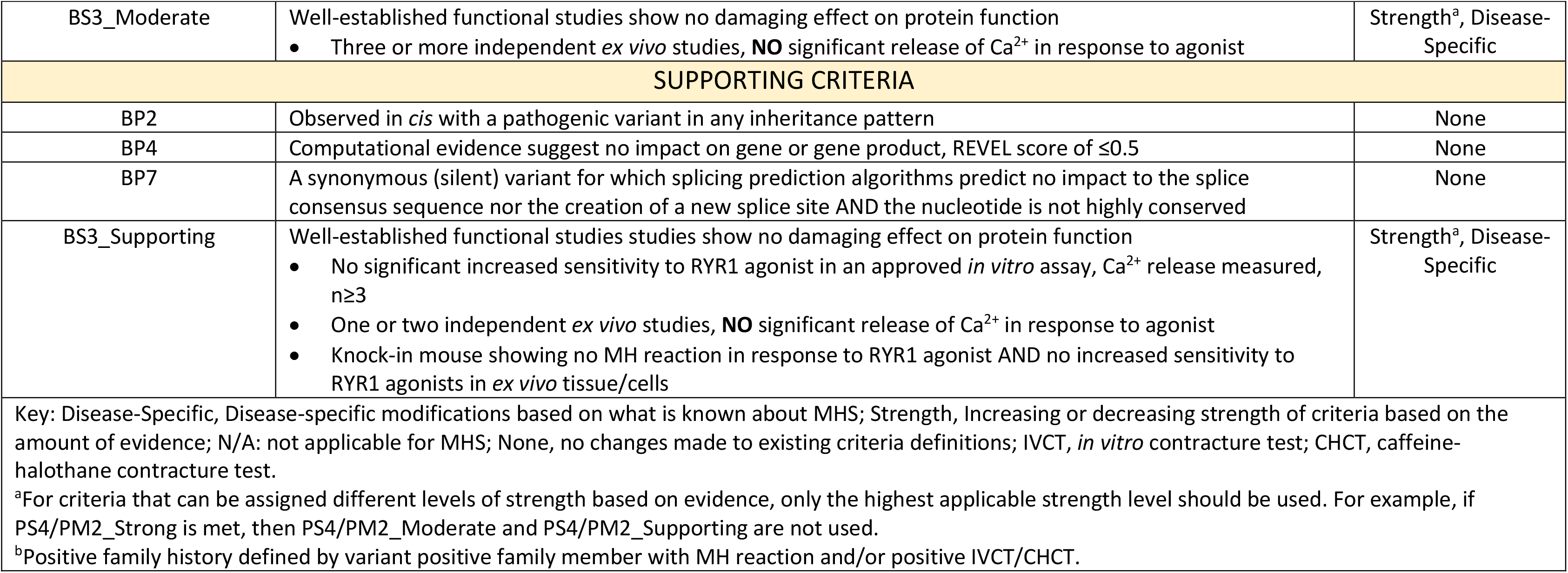
Modified ACMG criteria suggested for autosomal dominantly inherited RYR1/MH.

| Criteria | Criteria Description | Specification |
| --- | --- | --- |
| VERY STRONG CRITERIA |  |  |
| PS2/PM6_Very Strong | Each proven *de novo* case, 2 points, each assumed *de novo* case, 1 point, ≥8 points | Strength<sup>a</sup> |
| STRONG CRITERIA |  |  |
| PS1 | Same amino acid change as a previously established pathogenic variant regardless of nucleotide change<br>Previously established pathogenic variant must reach a classification of pathogenic without PS1 | None |
| PS2/PM6_Strong | Each proven *de novo* case, 2 points, each assumed *de novo* case, 1 point, a total of 4-7 points | Strength<sup>a</sup> |
| PS3 | Well-established functional studies supportive of a damaging effect on protein function<br>Knock-in mouse showing MH reaction in response to RYR1 agonist AND increased sensitivity to RYR1 agonists in *ex vivo* tissue/cells | Strength<sup>a</sup>, Disease-Specific |
| PS4 | The prevalence of the variant in affected individuals significantly increased compared with the prevalence in controls<br>≥7 MH case points. Proband with a personal or family history of an MH event are awarded 0.5 points, probands with a personal or family history<sup>b</sup> of a positive (MHS) IVCT/CHCT are awarded an additional 0.5 points. Popmax in gnomAD ≤0.00006<br>For variants with popmax MAF gnomAD >0.00006, an odds ratio of ≥18.7 when comparing MH case points to allele count in gnomAD can qualify. Popmax in gnomAD must be <0.0038 | Strength<sup>a</sup>, Disease-Specific |
| PP1_Strong | Co-segregation with disease in ≥7 reported meioses | Strength<sup>a</sup> |
| MODERATE CRITERIA |  |  |
| PM1 | Located in a mutational hot spot and/or critical and well established functional domain<br>Residues 1-552 (N-terminal region) and 2,101-2,458 (central region) | Disease-Specific |
| PM5 | Missense change at an amino acid residue where a different missense variant previously determined to be pathogenic<br>Previously established pathogenic variant must reach a classification of pathogenicity without PM5<br>Grantham score for alternate pathogenic variant must be less than for variant being assessed | None |
| PS2/PM6_Moderate | Each proven *de novo* case, 2 points, each assumed *de novo* case, 1 point, a total of 2-3 points | Strength<sup>a</sup> |
| PS3_Moderate | Well-established functional studies supportive of a damaging effect on protein function<br>Increased sensitivity to RYR1 agonist in HEK293 *in vitro* assay, Ca<sup>2+</sup> release significantly increased compared to WT, controls to include known pathogenic and benign variants, n≥3.<br>Three or more independent *ex vivo* studies all showing release of Ca<sup>2+</sup> in response to RYR1 agonist<br>Knock-in mouse showing MH reaction in response to RYR1 agonist OR increased sensitivity to RYR1 agonists in *ex vivo* tissue/cells (but not both, which would be PS3_strong) | Strength<sup>a</sup>, Disease-Specific |
| PS4_Moderate | The prevalence of the variant in affected individuals is significantly increased compared with the prevalence in controls | Strength<sup>a</sup>, Disease-Specific |
|  | 2-6 MH case points. Probands with a personal or family history<sup>b</sup> of an MH event are awarded 0.5 points, probands with a personal or family history of a positive (MHS) IVCT/CHCT are awarded an additional 0.5 points. Popmax in gnomAD $\leq 0.00006$<br>For variants with popmax MAF in gnomAD $> 0.00006$, an odds ratio of $\geq 4.33$ when comparing MH case points to allele count in gnomAD can qualify. Popmax in gnomAD must be $< 0.0038$ |  |
| PP1_Moderate | Co-segregation with disease in 5-6 reported meioses | Strength<sup>a</sup> |
| PP3_Moderate | Multiple lines of computational evidence support a deleterious effect on the gene or gene product<br>Use REVEL score of $\geq 0.85$ | Strength<sup>a</sup> |
| SUPPORTING CRITERIA |  |  |
| PP1 | Co-segregation with disease in 3-4 reported meioses | Strength<sup>a</sup> |
| PS2/PM6_Supporting | Each proven *de novo* case, 2 points, each assumed *de novo* case, 1 point, a total of 1 point | Strength<sup>a</sup> |
| PS3_Supporting | Well-established functional studies supportive of a damaging effect on protein function<br>Two independent *ex vivo* studies all showing release of $\text{Ca}^{2+}$ in response to RYR1 agonist | Strength<sup>a</sup>, Disease-Specific |
| PS4_Supporting | The prevalence of the variant in affected individuals is significantly increased compared with the prevalence in controls<br>1 MH case point. Probands with a personal or family history<sup>b</sup> of an MH event are awarded 0.5 points, probands with a personal or family history of a positive (MHS) IVCT/CHCT are awarded an additional 0.5 points. Popmax in gnomAD $\leq 0.00006$<br>For variants with popmax MAF in gnomAD $> 0.00006$, an odds ratio of $\geq 2.08$ when comparing MH case points to allele count in gnomAD can qualify. Popmax in gnomAD must be $< 0.0038$ | Strength<sup>a</sup>, Disease-Specific |
| PM1_Supporting | Located in a mutational hot spot and/or critical and well established functional domain<br>Residues 4,631-4,991 (C-terminal region) | Strength, Disease-Specific |
| BENIGN CRITERIA |  |  |
| STAND ALONE CRITERIA |  |  |
| BA1 | Popmax allele frequency $> 0.0038$ (0.38%) | Disease-Specific |
| STRONG CRITERIA |  |  |
| BS1 | Popmax allele frequency $> 0.0008$ (0.08%) | Disease-Specific |
| BS2 | Observed in a healthy adult individual for a recessive (homozygous), dominant (heterozygous), or X-linked (hemizygous) disorder with full penetrance expected at an early age.<br>Two or more variant positive individuals with a negative IVCT/CHCT test | Disease-Specific |
| MODERATE CRITERIA |  |  |
| BS2_Moderate | Observed in a healthy adult individual for a recessive (homozygous), dominant (heterozygous), or X-linked (hemizygous) disorder with full penetrance expected at an early age.<br>One variant positive individual with a negative IVCT/CHCT test | Strength<sup>a</sup>, Disease-Specific |
| BS3_Moderate | Well-established functional studies show no damaging effect on protein function<br>Three or more independent *ex vivo* studies, **NO** significant release of Ca<sup>2+</sup> in response to agonist | Strength<sup>a</sup>, Disease-Specific |
| SUPPORTING CRITERIA |  |  |
| BP2 | Observed in *cis* with a pathogenic variant in any inheritance pattern | None |
| BP4 | Computational evidence suggest no impact on gene or gene product, REVEL score of ≤0.5 | None |
| BP7 | A synonymous (silent) variant for which splicing prediction algorithms predict no impact to the splice consensus sequence nor the creation of a new splice site AND the nucleotide is not highly conserved | None |
| BS3_Supporting | Well-established functional studies studies show no damaging effect on protein function<br>No significant increased sensitivity to RYR1 agonist in an approved *in vitro* assay, Ca<sup>2+</sup> release measured, n≥3<br>One or two independent *ex vivo* studies, **NO** significant release of Ca<sup>2+</sup> in response to agonist<br>Knock-in mouse showing no MH reaction in response to RYR1 agonist AND no increased sensitivity to RYR1 agonists in *ex vivo* tissue/cells | Strength<sup>a</sup>, Disease-Specific |
| Key: Disease-Specific, Disease-specific modifications based on what is known about MHS; Strength, Increasing or decreasing strength of criteria based on the amount of evidence; N/A: not applicable for MHS; None, no changes made to existing criteria definitions; IVCT, *in vitro* contracture test; CHCT, caffeine-halothane contracture test.<br><sup>a</sup>For criteria that can be assigned different levels of strength based on evidence, only the highest applicable strength level should be used. For example, if PS4/PM2_Strong is met, then PS4/PM2_Moderate and PS4/PM2_Supporting are not used.<br><sup>b</sup>Positive family history defined by variant positive family member with MH reaction and/or positive IVCT/CHCT. |  |  |

### Criteria Dropped for MH/*RYR1*: PVS1/PM3/PM4/PP2/PP4/BS4/BP1/BP3/BP5

These criteria were dropped based on the biology of MH/*RYR1*. See supplemental information for details.

### Criteria Used According to General Guidelines: PS1/PS2/PM5/PM6/PP1/BP2/BP7

These criteria were retained in the *RYR1*/MH-specific guidelines including adaptations as recommended by the Clingen Sequence Variant Interpretation (SVI) committee (PS2/PM6, weighting of *de novo* observations, https://clinicalgenome.org/site/assets/files/3461/svi_proposal_for_de_novo_criteria_v1_0.pdf) and the Cardiomyopathy EP (PP1, weighting segregation events in families).^14^ We made further modifications to the ACMG/AMP criteria, which may not be specific to *RYR1*/MH. The PS1 (same amino acid change, different nucleotide change) and PM5 (different amino acid change, same codon) criteria were modified such that in order to use either of them, one variant should reach an assessment of pathogenic based on criteria other than PS1 and PM5. Then, PS1 or PM5 may be used for a second variant. Furthermore, for PM5, we added a requirement that the Grantham score difference compared to reference of the new variant must be greater than that for the previously identified pathogenic variant compared to reference. For criterion BP2 (evidence against pathogenicity based on presence of known pathogenic variant) it is suggested that only variants identified in *cis* with the variant under review be considered. Because the occurrence of biallelic pathogenic *RYR1* variants has been described in MHS,^3,15^ two variants in *trans* is not considered evidence against pathogenicity for *RYR1*/MH. Finally, BP7 concerns synonymous variants without predicted effects on splicing. As *RYR1*/MH primarily results from missense alterations, BP7 is used as recommended.

### Criteria Specified for RYR1/MH: BA1/BS1/PS4/PM2/BS2/PS3/BS3/PM1/PP3/BP4 Allele Frequency Specificiations: BA1/BS1/PS4/PM2

BA1 and BS1 use minor allele frequencies (MAF) in population datasets to support benign classification for common variants. The BA1 criterion is considered stand alone and was originally set to 0.05 (5%) MAF.^16^ It has been suggested that BA1 can be defined as the combined MAF for all pathogenic variants in the population for the gene/disease dyad with the understanding that any one variant should have a lower MAF than the combined total. To determine a gene/disease-specific cutoff for BA1, disease prevalence, penetrance, and gene contribution need to be considered. This can be estimated by the formula: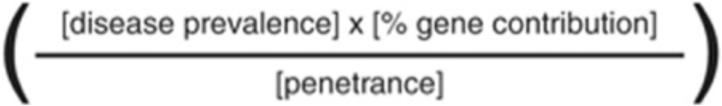.^14^ The prevalence of MH (defining the disorder as MH, not MHS) in the population can be estimated using the frequency of MH events in individuals exposed to triggering agents. The frequency of events is as high as 1/10,000 pediatric anesthesias.^2^ The rate of adult MH events seems lower than that of children^17^ but the underlying genetic risk is assumed to be the same. The gene contribution of RYR1 to MH is ~76% depending on ethnicity.^7^ Calculating thresholds for BA1 relies on an accurate estimate of penetrance, which is very difficult to determine for MHS.^18^ In lieu of using an estimate for MHS penetrance, we instead substituted a value of 1%, as it is a reasonable boundary between the penetrance of a Mendelian disorder variant and that of a risk allele. This value is nearly certain to be lower than the actual penetrance of MHS, but underestimating this value is conservative with respect to the outcome in that it will numerically raise BA1, which would lead to fewer variants being classified as benign based on this single criterion. Using 0.01 to adjust our calculated BA1 allows for a BA1 MAF of 0.0038 (0.38%).

In addition to a stand alone MAF (BA1), BS1 defines the MAF at which a variant is considered to have strong evidence against pathogenicity. The field has been moving to define BS1 based on the contribution of the most common pathogenic allele for a disorder. For *RYR1*/MH, we calculated BS1 considering the frequency of MH reactions in children (1/10,000) a value of 0.01 substituted for penetrance (as explained above), and a maximum individual allele contribution of 16%.^7^ Correcting for alleles/person gives a BS1 value of 0.0008 (0.08%).

While a high MAF of a variant in controls can be used to refute pathogenicity, criterion PM2 gives weight for absence or very low frequency of a variant in control populations. Based on observations that the majority of possible *RYR1* missense variants (~30,000 variants) are not represented in gnomAD (2,800 *RYR1* missense variants) and many known pathogenic variants (assessed without the use of PM2) are present in gnomAD, it is unlikely that the absence of a variant in gnomAD is support for pathogenicity. While the absence or low frequency of a variant in gnomAD has little value alone, it is an important component of weighting the presence of a variant in affected individuals. PS4 takes into consideration the prevalence of the variant in affected individuals compared to controls. For *RYR1*/MH, we modified the PS4 criterion using a point system, awarding 0.5 case points for each unrelated proband reported to have undergone an MH event and awarding an additional 0.5 case points for a positive *in vitro* contracture (IVCT) or caffeine-halthane contracture (CHCT) diagnostic test in either the proband or a variant-positive family member. The strength level of PS4 is based on odds ratios comparing total case points, an approximation of the total number of cases of MH investigated in the literature (3,000) and a MAF of 0.00006 for an allele with high coverage in the NFE population (approximately 7/113,000 alleles). When popmax frequency in gnomAD is >0.00006, cases can be counted and compared to alleles in the gnomAD population with the highest MAF by calculating an odds ratio (OR, MedCalcs online calculator (https://www.medcalc.org/calc/odds_ratio.php). Strength levels are awarded according to the following system: PS4 for ≥7 MH case points or an OR of 18.7; PS4_Mod for 2-6 MH cases points or an OR of 4.33; and PS4_Sup for one MH case point or an OR of 2.08. Every effort needs to be made to avoid double counting of cases reported in multiple studies. The Bayesian framework for the classification of variants using the ACMG/AMP criteria was used to set the OR value for each strength level.^13^

### Disease-Specific Phenotype: BS2

The IVCT/CHCT diagnostic tests have low false negative rates^19,20^ and can be used to determine MHS status in individuals who carry *RYR1* variants. A negative IVCT or CHCT result supports benign status. Two or more unrelated individuals with a negative result allow BS2 to be applied. One individual with a negative result allows BS2_Mod.

### Functional Assay Specifications: PS3/BS3

Functional characterization is a crucial determinant of the pathogenicity of *RYR1* variants in the MH field.^21^ Within the ACMG/AMP guidelines, functional assay results are used for PS3 (well-established *in vitro* or *in vivo* functional studies supportive of a damaging effect) and BS3 (well-established *in vitro* or *in vivo* functional studies show no damaging effect on protein function or splicing). RYR1 is a homotetrameric calcium channel in the sarcoplasmic reticulum (SR) of skeletal muscle important in excitation-contraction coupling. Volatile anesthetics and depolarizing muscle relaxants can cause increased release of SR calcium in a dysfunctional RYR1 channel resulting in MH. When considering functional assays for variant assessment it is desirable to identify assays that are closely related to the physiologic defect causative of disease. For *RYR1*/MH, assays that measure release of calcium in response to pharmacologic agents are considered good representations of the disease mechanism. Well-recognized assays include transfection of *RYR1* cDNA into either HEK293 cells, CHO cells, or *RYR1* knockout myotubes (dyspedic) followed by SR calcium release measurement in response to caffeine, halothane, voltage/potassium, or 4-chloro-*m*-cresol (4-C*m*C). A significant decrease in the EC_50_ for the sensitivity of calcium release compared to wildtype RYR1, is considered evidence for pathogenicity. Multiple replicates for each variant within a single instance of the assay are necessary to determine significance of these values. Positive (pathogenic) and negative (benign) controls support that the assay categorizes the variants accurately. For the purpose of assessing *RYR1* transfection studies to weight PS3, results are dichotomized into pathogenic EC_50_ values that are significantly decreased as compared to WT versus benign EC_50_ values that are not significantly decreased. For *RYR1* pathogenicity assessment, the whole of prior published work (Figure 1, Table S2)^22^ allows us to consider transfection assays in HEK293 cells using photometry/imaging to measure calcium release a well defined functional test. However, recommendations for increased stringency in analyses of functional data have recently been suggested.^23^ To determine the appropriate PS3 weight based on HEK293 transfection assays we have considered results published in the literature including results for a total of 35 variants assessed to be likely pathogenic or pathogenic (LP/P) without the use of functional data, and ten control variants including variants associated with CCD (8) and common variants (2). Of the 35 LP/P variants, 29 have been shown to reduce the calcium release EC_50_ in response to RYR1 agonsits. Five variants have shown discordant results across assays, and one variant has shown an EC_50_ increase. Of the ten control variants, one variant has shown an EC_50_ reduction in response to agonist and nine variants have either shown no response to agonist (6) or a response similar to WT RYR1 (3). This set of variants suggests a likelihood ratio for an EC_50_ reduction of 9.11:1 with a 95% confidence interval of 1.4:1 to 59:1. This level of support is above the threshold for moderate evidence (4.33:1 odds). We suggest that functional evidence supporting pathogenicity from HEK293 cells be used at the level of moderate. When the field generates additional data for control variants the weighting of PS3 for this assay should be reconsidered.

**Figure 1.**
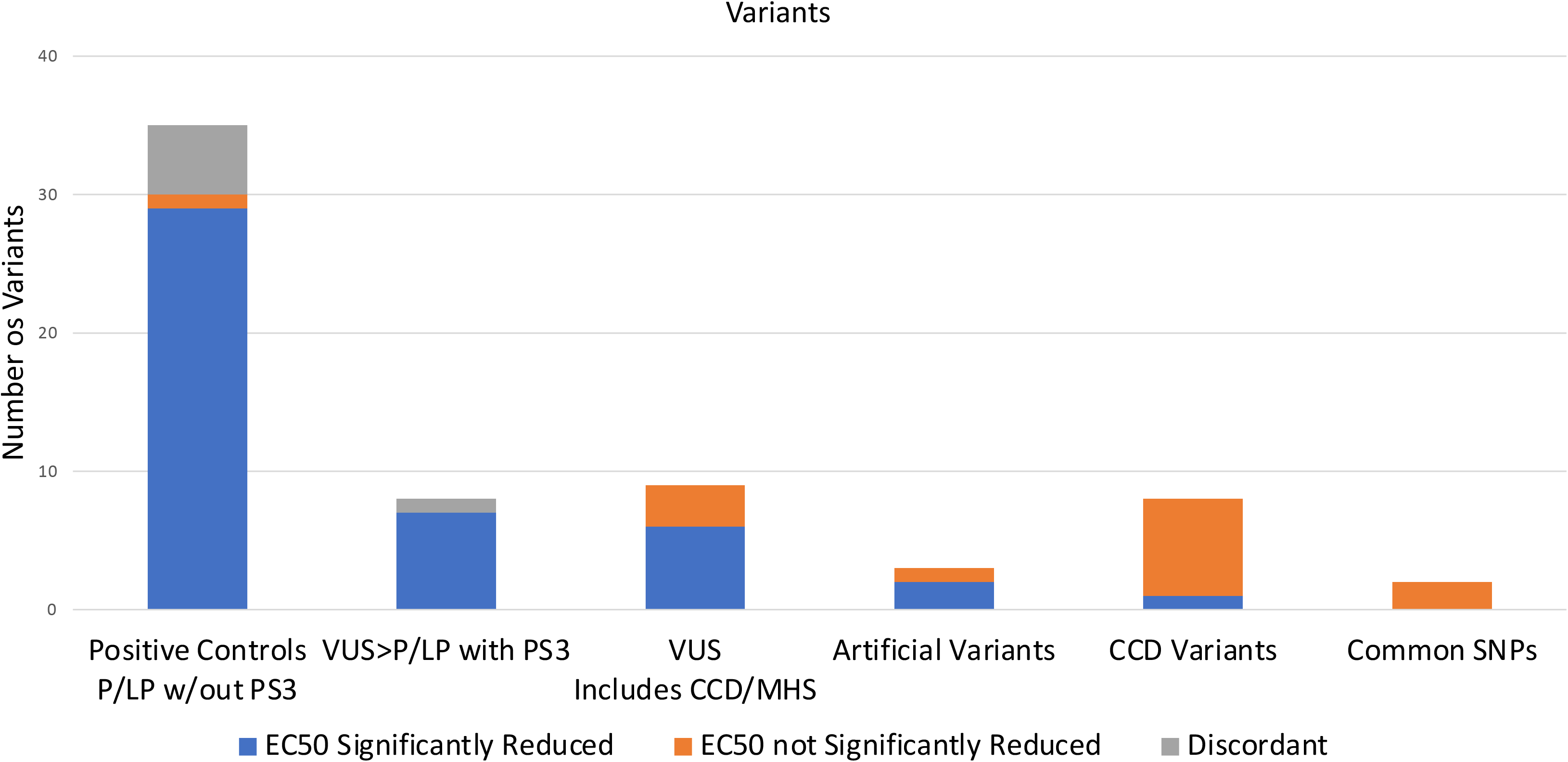
Cumulative HEK293 transfection assay data for RYR1 variants from the literature. Variants are grouped according to pathogenicity assessment without consideration of PS3/BS3 (functional data).

**Figure 2.**
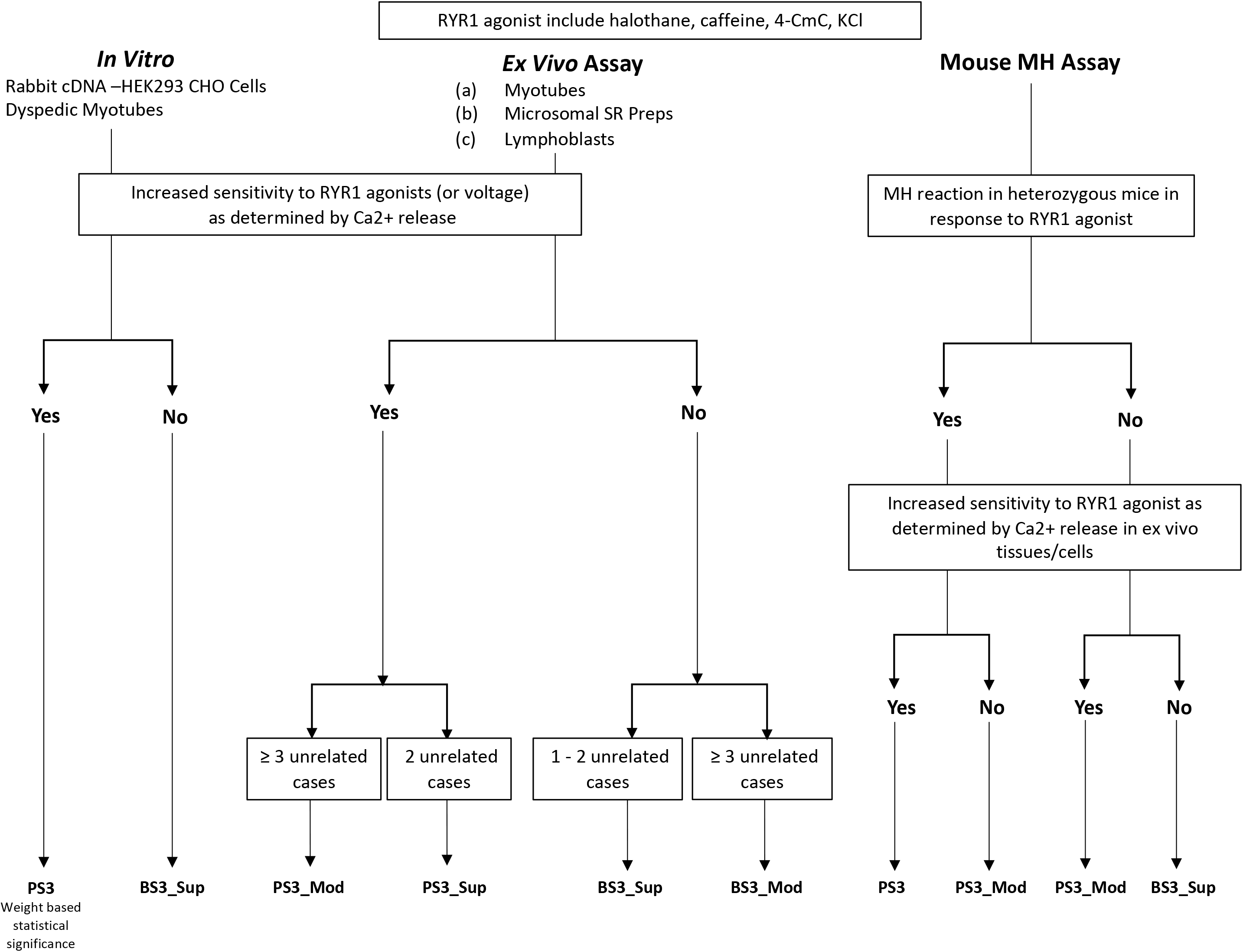
Decision tree for weighting functional evidence PS3/BS3.

While positive evidence (reduced EC_50_) is considered moderate support for pathogenicity, reduced penetrance and the limitations of expression systems,^24^ suggest a non-significant change in EC_50_ values may not support benign status at a moderate level. For that reason it was decided that lack of response to agonists be weighted as supporting evidence, BS3_Sup. Regarding other *in vitro* assays that test calcium release in response to agonists, where historical data were limited, we suggest that multiple controls be run in parallel and statistical analysis be used to determine the level of strength for PS3 according to the Bayesian framework. Future data from control variants will allow for reconsideration of increased weight of PS3.

In addition to *in vitro* assays, the *RYR1*/MH field has established *ex vivo* assays measuring calcium release in patient cells. These assays do not isolate the *RYR1* variant from other potential variants (in *RYR1*, *CACNA1S*, or other MHS-associated genes), which may affect calcium release. Rather, these assays are a measure of the cellular phenotype in the patient. Although we recognize this limitation of *ex vivo* studies, we also recognize that they have utility. As the main concern for such assays is the potential presence of other variants, this concern is mitigated if multiple unrelated individuals with the same primary variant are shown to exhibit enhanced *ex vivo* sensitivity to agonist. Two unrelated individuals with *ex vivo* tests showing increased sensitivity of calcium release in response to agonist allow PS3_Sup. For variants where ≥3 unrelated individuals had *ex vivo* tests showing increased sensitivity of calcium release, PS3_Mod can be applied. *Ex vivo* tests that do not show increased sensitivity of calcium release in response to agonist (negative result) support a benign status of the variant. BS3_Sup can be applied if one or two unrelated individuals are tested with negative results, when ≥3 unrelated individuals are tested and all results are negative BS3_Mod can be applied. Knock-in mouse models created to date to test *RYR1* variants have shown MH reactions in response to exposure to volatile anesthetic and *ex vivo* studies of muscle samples from these mice show increased ligand sensitivity of calcium release as compared to WT.^25–28^ When knock-in mice have an MH reaction in response to agonist, and where *ex vivo* studies show increased calcium release as compared to WT in response to agonist, PS3 can be awarded. For mouse models where either an MH crisis can be triggered by agonist or *ex vivo* assays show increased calcium release, but both conditions are not met, PS3_Mod is awarded. For mouse models that do not exhibit an MH reaction when exposed to agonist and *ex vivo* studies do not show increased release of calcium, BS3_Sup is can be awarded.

### Hotspot Specifications: PM1

The ACMG/AMP criteria includes moderate weight for variation in critical protein domains or mutational hotspots, PM1. While critical domains may be well-defined for a protein, the concept of mutational hotspot is less clearly defined in the field. A general rule for consideration of a mutational hotspot would be an excess of pathogenic variation as compared to benign variation. In MH, variants have been noted to cluster in three regions of *RYR1* identified as “hotspots” historically: the N-terminal region (residues 1-552), the central region (residues 2,101-2,458) and the C-terminal region (4,631-4,991).^29^ Rather than defining clear functional domains, these regions are defined by an increase in variation identified in individuals with MH. We assessed this criterion using a test set of 21 variants (Table S3) assessed to be pathogenic for MH without the use of PM1 and 27 benign variants (Table S4) that met criteria BA1. This set of variants suggests a likelihood ratio for hotspots of 24:1 with a 95% confidence interval of 3.5:1 to 166:1 (Table 2). This level of support is above the threshold for strong evidence (18.7:1 odds) and the lower bound of that confidence interval is above supporting (2.1:1). This would suggest that PM1 could be modified to PM1_strong. However, because there is a significant bias in the literature toward identifying pathogenic variants in the hotspots, to avoid the possibility of overestimating pathogenicity, we suggest instead using PM1 at its default level of moderate for variants in the N-terminal and central regions. We suggest using PM1 at a supporting level for variants in the C-terminal region as variants in this region may be associated with CCD and not cause MH. Future studies that interrogate the gene without these biases should provide additional data on the positional skewing of pathogenic variants, which could allow us to upgrade this to strong in the future.

**Table 2.** Distribution of 21 pathogenic and 27 benign variants in relation to position of defined *RYR1*/MH hotspots. Likelihood ratios calculated based on distribution.

| Presence in HotSpot | Pathogenic | Benign | Likelihood ratio (LR) | Inverse LR | 95% CI |
| --- | --- | --- | --- | --- | --- |
| HotSpot<br>(1-552; 2101-2458;<br>4631-4991) | 18 | 1<sup>a</sup> | 24.00 |  | 3.48-165.77 |
| Non-HotSpot | 3 | 27 | 0.148 | 6.76 | 0.05-0.42 |

<sup>a</sup>No benign variants were identified in the hotspot regions, for calculation of LR we used a value of 1.

### Computational Evidence: PP3/BP4

The PP3 and BP4 criteria consider computational evidence estimating the impact of a variant on protein function. REVEL is an ensemble method based on a number of individual tools and precomputed scores are available for all missense variants (https://omictools.com/revel-tool).^12^ Importantly, REVEL does not consider population frequency, which reduces double counting of evidence. Using a set of 22 pathogenic variants determined to be pathogenic without the use of PP3 the and 27 benign variants described above, we tested the likelihood ratios of the predictive power of REVEL in several iterations. We settled on a trichotomization of scores with PP3, (computational evidence supporting pathogenicity), requiring a REVEL score of ≥0.85 and BP4, (computational evidence against pathogenicity), requiring a REVEL score of ≤0.5 (Table 3). These results suggest that PP3 and BP4 could be employed at the strong level. We chose to reduce PP3 to moderate as it was close to the Bayesian strong cutoff of 18.7:1 odds.^13^ Based on piloting these criteria it was determined that BP4 should be used at supporting and only implemented with other criteria. Using the Bayesian framework, BP4 in isolation results in an assessment of likely benign (LB) and it was determined that additional evidence should be available for a LB classification. For a fuller explanation of deriving such likelihood ratios, see Supplemental information.

**Table 3.** REVEL score distribution for 22 pathogenic and 27 benign variants for *RYR1*/MH. Likelihood ratio for separation of pathogenic and benign variants based on REVEL scores using cutoff values of >0.85 and <0.5.

| REVEL score | Pathogenic | Benign | Likelihood ratio (LR) | Inverse LR | 95% CI |
| --- | --- | --- | --- | --- | --- |
| >0.85 | 19 | 1<sup>a</sup> | 23.13 |  | 3.35-159.96 |
| 0.5-0.85 | 3 | 8 | 0.46 | 2.19 | 0.14-1.53 |
| <0.5 | 1<sup>a</sup> | 19 | 0.06 | 15.63 | 0.01-0.44 |

<sup>a</sup>No benign variants were identified with a REVEL score >0.85 and no pathogenic variants were identified with a REVEL score <0.5, for calculation of LR we used a value of 1.

### Piloting RYR1/MH Assessment Criteria

We applied these modified criteria to 44 variants EMHG determined to be “diagnostic mutations” and 40 *RYR1* variants with pathogenicity assessments for MH in ClinVar. The classification of each of the variants is shown in Table S3 and Table S5. Four variants included in the EMHG variant list (https://www.emhg.org/diagnostic-mutations) were excluded from this assessment as they are not associated with MH. Of the remaining 44 EMHG variants, we assessed 30 to be pathogenic (P), 12 to be likely pathogenic (LP), and two to be variants of uncertain significance (VUS). Variant c.1589G>A p.(Arg530His) was assessed as VUS and had limited functional data including a single *ex vivo* sample^30^, which did not meet PS3_Sup based on the requirement for a minimum of two unrelated individuals. Variant c.1598G>A p.(Arg533His) was assessed as VUS based on functional data (PS3_Mod) and presence in a hotspot (PM1). PS4 was not met by this variant based on a high allele count (32 alleles) in gnomAD.

The revised criteria were applied to 40 additional variants with pathogenicity assessments for MH in ClinVar. Ten variants had conflicting pathogenicity assessments for MH (pathogenicity assessments not indicated for MH were not considered), nine B/LB/VUS and one P/LP/VUS. Five variants with B/LB/VUS assessments in ClinVar were determined to be B/LB based on BA1/BS1. The remaining five discordant variants were assessed to be VUS. Of the remaining 30 variants, 14 were reported as P/LP, 11 as B/LB and five as VUS. Applying the revised ACMG criteria 12/14 variants with an assessment of P/LP in ClinVar and 3/11 variants with an assessment of B/LB in ClinVar were assessed as VUS. All variants assessed as B/LB (13) using our criteria had ether BA1 or BS1 applied. The 19/24 variants assessed as VUS had limited data, only five VUS variants had data that refuted pathogenicity (5/24, 21%).

## CONCLUSIONS

As an expert panel within ClinGen, we set out to adapt the ACMG pathogenicity criteria for assessment of *RYR1* variants as related to autosomal dominanty inherited MH. Combining expertise of anesthesiologists, physiologists, biochemists, and geneticists allowed for a thorough evaluation of factors that should be considered. It is also important to recognize that we successfully unified the efforts of the American-based ACMG/AMP criteria with the extensive expertise and experience of the European Malignant Hyperthermia Group, benefiting from both. In revising these guidelines, we have considered the statistical evidence weight as it relates to the Bayesian adaptation of the ACMG scoring system. Weighting of evidence using statistical measures should allow for a more robust and consistent pathogenicity assessment framework. The revised *RYR1*/MHS specific criteria should allow clinical laboratories to more consistently assess these variants based on expert guidelines. These recommendations should be especially useful to laboratories that interpret *RYR1* variants as secondary findings in exome and genome sequencing. That MH is a pharmacogenetic trait with relatively low penetrance makes it especially challenging to interpret for laboratories that do not peform a high volume of diagnostic *RYR1* testing. The availability of these three star ClinGen interpretations should significantly reduce the amount of time that these secondary findings evaluations consume and should increase the consistency of the interpretations, as has been demonstrated for the generic ACMG/AMP pathogenicity recommendations.^31^ As well, the *RYR1*/MH expert panel will continue to curate variants and deposit assessments into ClinVar. Standardized ClinVar assessments will be useful to laboratories identifying variants as secondary findings that may not have *RYR1*/MH expertise. Moving forward, the field should strive to increase relevant data through functional studies and shared case documentation allowing variants to move from an assessment of VUS to either LB/B or LP/P. Beyond secondary findings, ClinGen interpretations of *RYR1* variant pathogenicity will allow the field to consider pre-surgicial screening of patients toward elimination of MH morbidity and mortality.^32^

## ACKNOWLEDGEMENTS

ClinGen is primarily funded by the National Human Genome Research Institute (NHGRI), through the following three grants: U41HG006834, U41HG009649, U41HG009650. ClinGen also receives support for content curation from the Eunice Kennedy Shriver National Institute of Child Health and Human Development (NICHD), through the following three grants: U24HD093483, U24HD093486, U24HD093487. The content is solely the responsibility of the authors and does not necessarily represent the official views of the National Institutes of Health. JJJ and LGB were supported by NIH grant HG200359-12.

RTD is supported by grant R01 AR053349. PH is supported by grants from the NIH: National Institute of Arthritis, Musculoskeletal and Skin Diseases (2P01 AR-05235, 1R01AR068897-01A1). SR is funded by merit award from the Department of Anesthesia and Pain Medicine, University of Toronto, Canada.

## AUTHOR CONTRIBUTION

Conceptualization: L.G.B.; Data curation: J.J.J., S.G.G., K.S., L.G.B.; Formal Analysis: J.J.J., L.G.B.; Methodology: J.J.J., R.T.D., T.G., S.G.G., P.M.H., S.R., L.A.S., N.S., R.S., K.S., J.W., N.R., L.G.B.; Project Administration: J.J.J., L.G.B.; Writing – original draft: J.J.J., L.G.B.; Writing – review & editing: R.T.D., T.G., S.G.G., P.M.H., S.R., L.A.S., N.S., R.S., K.S., J.W., H.R.

