## Supplemental Information for "Variant Curation Expert Panel Recommendations for RYR1 Pathogenicity Assertions in Malignant Hyperthermia Susceptibility"

Table S1. Explanation of modified ACMG criteria for autosomal dominantly inherited *RYR1*/MH.

**VERY STRONG EVIDENCE OF PATHOGENICITY**

| | |
|---|---|
| **PVS1** | Null variant (nonsense, frameshift, canonical +/- 1 or 2 splice sites, initiation codon, single or multi-exon deletion) in a gene where loss of function (LOF) is a known mechanism of disease.<br>**MHS-*RYR1*:** PVS1 is not applicable. MHS is due to gain of function variants in *RYR1*. |
| **PS2/PM6_ Very Strong** | *De novo* in a patient with the disease and no family history. Counts BOTH proven and unproven *de novo* cases.<br>Note: Confirmation of paternity only is insufficient. Egg donation, surrogate motherhood, errors in embryo transfer, etc. can contribute to non-maternity.<br>**MHS-*RYR1*:** PS2/PM6 follow SVI recommendation for *de novo* criteria. Each proven de *novo* case gets 2 points, each unproven *de novo* case gets 1 point, PS2/PM6_Very Strong applied if ≥8 points.<br><br>Note: The family history should be negative for MH events, central core disease, or exertional heat related illness. |

**STRONG EVIDENCE OF PATHOGENICITY**

| | |
|---|---|
| **PS1** | Same amino acid change as a previously established pathogenic variant regardless of nucleotide change.<br>**MHS-*RYR1*:** PS1 is applicable as described. As with PM5, the initial variant determined to be pathogenic must reach an assessment of pathogenic without using this criterion (no double counting). |
| **PS2/PM6_ Strong** | *De novo* in a patient with the disease and no family history. Counts BOTH proven and unproven *de novo* cases.<br>Note: Confirmation of paternity only is insufficient. Egg donation, surrogate motherhood, errors in embryo transfer, etc. can contribute to non-maternity.<br>**MHS-*RYR1*:** PS2/PM6 follow SVI recommendation for *de novo* criteria. Each proven *de novo* case gets 2 points, each unproven *de novo* case gets 1 point, PS2/PM6_Strong applied if 4-7 points.<br><br>Note: The family history should be negative for MH events, central core disease, or exertional heat related illness. |
| **PS3** | Well-established *in vitro* or *ex vivo* functional studies or knock-in mouse studies supportive of a damaging effect on the gene or gene product.<br>**MHS-*RYR1*:**<br>• *In vitro* assays showing increased sensitivity to RYR1 agonist (halothane, caffeine, 4-CmC, KCl, voltage) can be used for PS3. All assays require appropriate controls such that likelihood ratios are ≥18.7. Historical data, when available, can be used to validate the assay.<br>• MH reaction in response to RYR1 agonist in a knock-in mouse model, requires **BOTH** MH reaction in heterozygous animals AND increased sensitivity to RYR1 agonist (halothane, caffeine, 4-CmC, KCl, voltage) in an approved ex *vivo* assay using knock-in mouse tissues, $Ca^{2+}$ release measured by fluorescence. Appropriate controls should be included. |
| **PS4** | The prevalence of the variant in affected individuals is significantly increased compared to the prevalence in controls.<br>**MHS-*RYR1*:** True case control studies do not exist in the *RYR1* literature with controls known to be negative for MHS. A modified PS4 is used for *RYR1* using MH case reports and data from gnomAD (Richards et al. Note 2).<br>• PS4_Strong requires ≥7 MH case points, one point is awarded for a proband with a personal or family history (in a variant positive individual) of an MH event AND a positive IVCT or CHCT diagnostic test (MHS), 0.5 points are awarded for a proband with a reported MH event but without an IVCT or CHCT diagnostic test. Popmax MAF in gnomAD ≤0.00006. |

| | |
|---|---|
| | • For variants with popmax MAF in gnomAD ≥0.00006, and below BA1 cutoff of 0.0038, MedCalcs online calculator can be used to calculate the OR using case points from the literature, an approximation of 3,000 cases (6,000 alleles) reported in the literature and allele counts from gnomAD (MedCalc; https://www.medcalc.net/statisticaltests/odds_ratio.php). An OR of ≥18.7 is required for PS4_Strong. |
| **PP1_Strong** | Co-segregation with disease in multiple affected family members.<br>**MHS-*RYR1*:** ≥7 meioses, only consider phenotype positive/variant positive individuals. To use PP1, no phenotype positive/variant negative individuals can be identified in a pedigree. |

**MODERATE EVIDENCE OF PATHOGENICITY**

| | |
|---|---|
| **PM1** | Located in a mutational hot spot and/or critical and well-established functional domain (e.g., active site of an enzyme) without benign variation.<br>**MSH-*RYR1*:** Residue regions: 1-552 (N-terminal region) and 2,101-2,458 (central region) are thought to be critical functional domains for MHS. |
| **PM2** | Absent from controls (or at extremely low frequency if recessive) in Exome Sequencing Project, 1000 Genomes or ExAC.<br>**MHS-*RYR1*:** PM2 is not used alone. |
| **PM3** | For recessive disorders, detected in trans with a pathogenic variant<br>Note: This requires testing of parents (or offspring) to determine phase.<br>**MHS-*RYR1*:** PM3 is not applicable. MHS is inherited as an autosomal dominant trait with reduced penetrance. |
| **PM4** | Protein length changes due to in-frame deletions/insertions in a non-repeat region or stop-loss variants.<br>**MHS-*RYR1*:** PM4 is not applicable. The majority of *RYR1* variants that are causative for MHS are missense variants. |
| **PM5** | Missense change at an amino acid residue where a different missense change determined to be pathogenic has been seen before.<br>**MHS-*RYR1*:** PM5 is applicable as described. As with PS1, the initial variant determined to be pathogenic must reach an assessment of pathogenic without using this criterion (no double counting). As well, the Grantham score difference for the initial variant determined to be pathogenic must be less than the Grantham score difference for the variant currently being assessed. |
| **PS2/PM6_ Moderate** | *De novo* in a patient with the disease and no family history. Counts BOTH proven and unproven *de novo* cases.<br>Note: Confirmation of paternity only is insufficient. Egg donation, surrogate motherhood, errors in embryo transfer, etc. can contribute to non-maternity.<br>**MHS-*RYR1*:** PS2/PM6 follow SVI recommendation for *de novo* criteria. Each proven *de novo* case gets 2 points, each unproven *de novo* case gets 1 point, PS2/PM6_Moderate applied for 2-3 points.<br><br>Note: Family history needs to be negative for MH events, central core disease, or exertional heat related illness. |
| **PS3_Moderate** | Well-established *in vitro* or *ex vivo* functional studies or knock-in mouse studies supportive of a damaging effect on the gene or gene product.<br>**MHS-*RYR1*:**<br>• *In vitro* assays showing increased sensitivity to RYR1 agonist (halothane, caffeine, 4-CmC, KCl, voltage) can be used for PS3_Moderate. All assays require appropriate controls such that likelihood ratios are ≥4.3. Historical data, when available, can be used to validate the assay.<br>• Historical assay data for transfection studies in HEK293 cells supports using this assay at the moderate strength level. Result showing increased sensitivity to RYR1 agonist (halothane, caffeine, 4-CmC) supports pathogenicity. Result must show significant increase in $Ca^{2+}$ release for agonist concentration ($EC_{50}$). Controls must include wildtype *RYR1* and known pathogenic |

| | |
|---|---|
| | variants that reach an assessment of LP/P without consideration of PS3. Assays should be run in triplicate.<br>• Three or more independent *ex vivo* studies (tissues from unrelated individuals) all showing increased release of $Ca^{2+}$ in response to RYR1 agonist (halothane, caffeine, 4-CmC, KCl, voltage). $Ca^{2+}$ release measured by fluorescence. Appropriate controls included. Result must show significant increase in $Ca^{2+}$ release at decreased agonist concentration.<br> ○ Patient tissues considered useful for PS3 (and BS3) include patient myotubes, microsomal SR preps, and lymphoblasts.<br>• MH reaction in response to RYR1 agonist in a knock-in mouse model, requires MH reaction in heterozygous animals OR increased sensitivity to RYR1 agonist (halothane, caffeine, 4-CmC, KCl, voltage) in an approved *ex vivo* assay using knock-in mouse tissues, $Ca^{2+}$ release measured by fluorescence. Appropriate controls should be included. |
| **PS4 _<br><br>Moderate** | The prevalence of the variant in affected individuals is significantly increased compared to the prevalence in controls.<br>**MHS-*RYR1*:**True case control studies do not exist in the *RYR1* literature with controls known to be negative for MHS. A modified PS4 is used for *RYR1* using MH case reports and data from gnomAD (Richards et al. Note 2).<br>• PS4_Moderate requires 2-6 MH case points, one point is awarded for a proband with a personal or family history (in a variant positive individual) of an MH event AND a positive IVCT or CHCT diagnostic test (MHS), 0.5 points are awarded for a proband with a reported MH event but without an IVCT or CHCT diagnostic test. Popmax MAF in gnomAD ≤0.00006.<br>• For variants with popmax MAF in gnomAD ≥0.00006, and below BA1 cutoff of 0.0038, MedCalcs online calculator can be used to calculate the OR using case points from the literature, an approximation of 3,000 cases (6,000 alleles) reported in the literature and allele counts from gnomAD (MedCalc; https://www.medcalc.net/statisticaltests/odds_ratio.php). An OR of ≥4.33 is required for PS4_Moderate. |
| **PP1_Moderate** | Co-segregation with disease in multiple affected family members.<br>**MHS-*RYR1*:** 5-6 meioses, only consider phenotype positive/variant positive individuals. In order to use PP1 no phenotype positive/variant negative individuals can be identified in a pedigree. |
| **PP3_Moderate** | Multiple lines of computational evidence support a deleterious effect on the gene or gene product (conservation, evolutionary, splicing impact, etc.).<br>**MHS-*RYR1*:** REVEL score of ≥ 0.85 is considered evidence in support of pathogenicity. (PMID:27666373) |

**SUPPORTING EVIDENCE OF PATHOGENICITY**

| | |
|---|---|
| **PP1** | Co-segregation with disease in multiple affected family members.<br>**MHS-*RYR1*:** 3-4 meioses, only consider phenotype positive/variant positive individuals. In order to use PP1 no phenotype positive/variant negative individuals can be identified in a pedigree. |
| **PP2** | Missense variant in a gene that has a low rate of benign missense variation and where missense variants are a common mechanism of disease.<br>**MHS-*RYR1*:** PP2 is not applicable. *RYR1* does not appear to be constrained for missense variation with a z-score of 1.92 in gnomAD. |
| **PP3** | Multiple lines of computational evidence support a deleterious effect on the gene or gene product (conservation, evolutionary, splicing impact, etc.).<br>**MHS-*RYR1*:** Upgraded to PP3_Moderate. |
| **PP4** | Patient's phenotype or family history is highly specific for a disease with a single genetic etiology.<br>**MHS-*RYR1*:** PP4 is not applicable, variants in *CACNA1S* also result in MHS. |
| **PP5** | Reputable source recently reports variant as pathogenic but the evidence is not available to the laboratory to perform an independent evaluation.<br>**MHS-*RYR1*:** PP5 has been dropped from the ACMG framework for variant assessment. |
| **PS2/PM6_** | *De novo* in a patient with the disease and no family history. Counts BOTH proven and unproven *de novo* cases. |

| | |
|---|---|
| **Supporting** | Note: Confirmation of paternity only is insufficient. Egg donation, surrogate motherhood, errors in embryo transfer, etc. can contribute to non-maternity.<br>**MHS-*RYR1*:** PS2/PM6 follow SVI recommendation for *de novo* criteria. Each proven *de novo* case gets 2 points, each unproven *de novo* case gets 1 point, PS2/PM6_Supporting applied for 1 point.<br><br>Note: The family history should be negative for MH events, central core disease, or exertional heat related illness. |
| **PS3_Supporting** | Well-established *in vitro* or *ex vivo* functional studies or knock-in mouse studies supportive of a damaging effect on the gene or gene product.<br>**MHS-*RYR1*:**<br>• *In vitro* assays showing increased sensitivity to RYR1 agonist (halothane, caffeine, 4-CmC, KCl, voltage) can be used for PS3_Supporting. All assays require appropriate controls such that likelihood ratios are ≥2.08. Historical data, when available, can be used to validate the assay.<br>• Two independent *ex vivo* studies (tissues from unrelated individuals) all showing increased release of $Ca^{2+}$ in response to RYR1 agonist (halothane, caffeine, 4-CmC, KCl, voltage). $Ca^{2+}$ release measured by fluorescence. Appropriate controls included. Result must show significant increase in $Ca^{2+}$ release at decreased agonist concentration.<br> ○ Patient tissues considered useful for PS3 (and BS3) include patient myotubes, microsomal SR preps and lymphoblasts. |
| **PS4_Supporting** | The prevalence of the variant in affected individuals is significantly increased compared to the prevalence in controls.<br>**MHS-*RYR1*:** True case control studies do not exist in the *RYR1* literature with controls known to be negative for MHS. A modified PS4 is used for *RYR1* using MH case reports and data from gnomAD (Richards et al. Note 2).<br>• PS4_Supporting requires one MH case point, one point is awarded for a proband with a personal or family history (in a variant positive individual) of an MH event AND a positive IVCT or CHCT diagnostic test (MHS), 0.5 points are awarded for a proband with a reported MH event but without an IVCT or CHCT diagnostic test.). Popmax MAF in gnomAD ≤0.00006.<br>• For variants with popmax MAF in gnomAD ≥0.00006, and below BA1 cutoff of 0.0038, MedCalcs online calculator can be used to calculate the OR using case points from the literature, an approximation of 3,000 cases (6,000 alleles) reported in the literature and allele counts from gnomAD (MedCalc; https://www.medcalc.net/statisticaltests/odds_ratio.php). An OR of ≥2.08 is required for PS4_Supporting. |
| **PM1_Supporting** | Located in a mutational hot spot and/or critical and well-established functional domain (e.g., active site of an enzyme) without benign variation.<br>**MSH-*RYR1*:** Residue region: 4,631-4,991 (C-terminal region) is thought to be a critical functional domain for MHS. Variants in this domain have been identified in MH and CCD. |

**STAND ALONE EVIDENCE OF BENIGN IMPACT**

| | |
|---|---|
| BA1 | Allele frequency is above 5% in Exome Sequencing Project, 1000 Genomes, or ExAC.<br>**MHS_*RYR1*:** An allele frequency of ≥ 0.0038 (0.38%) is used as a cut off based on popmax MAF in gnomAD (outbred population).<br><br>Calculating a stand-alone filtering frequency for MHS-RYR1 is complicated as neither the frequency nor the penetrance of MHS is well understood. Based on reduced penetrance and the requirement of a triggering event the incidence of MH events is expected to be lower than the incidence of MHS. Studies have reported an incidence of MH events as low as 1 in 10,000 to 1 in 250,000 anesthesias (PMID: 26709912).<br><br>Variants in *RYR1* are reported to account for ~ 76% of cases (PMID: 30236257).<br><br>Penetrance for MH is not well understood, we instead substituted a value of 1%, as it is a reasonable boundary between the penetrance of a mendelian disorder variant and that of a risk allele.<br><br>Maximum Prevalence of MHS = Prevalence of MH events / Penetrance<br>(1 event/10,000 children) / (1 event/100 causative alleles )<br>1 causative allele/100 children<br><br>MHS prevalence (0.01) * *RYR1* contribution (0.76) * Allele conversion (0.5) = 0.0038 |

**STRONG EVIDENCE OF BENIGN IMPACT**

| | |
|---|---|
| **BS1** | Allele frequency is greater than expected for disorder.<br>**MHS-*RYR1*:** An MH all allele frequency of ≥ 0.0008 (0.08%) is used as a cut off for popmax MAF in gnomAD (outbred population). Disease prevalence as explained for BA1.<br><br>Disease prevalence (0.01) * Maximum single *RYR1* variant contribution (0.16) * Allele conversion (0.5) = 0.0008 |
| **BS2** | Observed in a healthy adult individual for a recessive (homozygous), dominant (heterozygous), or X-linked (hemizygous) disorder with full penetrance expected at an early age.<br>**MHS-*RYR1*:** The absence of an MH reaction in a healthy individual cannot be used for BS2 due to reduced penetrance. BS2 is applicable if two or more unrelated variant positive individuals have negative results for either the IVCT or CHCT. |
| **BS3** | Well-established *in vitro* or *ex vivo* functional studies or knock-in mouse studies show no damaging effect on protein function.<br>**MSH-RYR1:** BS3 downgraded to BS3_Supporting for all negative data. |
| **BS4** | Lack of segregation in affected members of a family<br>**MSH-*RYR1*:** BS4 is not applicable. Phenotype for MHS is routinely determined based on the vitro contraction test (IVCT) that has a false positive rate of approximately 6% (PP1) or the caffeine-halothane contracture test (CHCT). As the phenotype in individuals who have not experienced an MH crisis cannot be reliably determined BS4 is not utilized. |

**MODERATE EVIDENCE FOR BENIGN IMPACT**

| | |
|---|---|
| **BS2_Moderate** | Observed in a healthy adult individual for a recessive (homozygous), dominant (heterozygous), or X-linked (hemizygous) disorder with full penetrance expected at an early age.<br>**MHS-*RYR1*:** The absence of an MH reaction in a healthy individual cannot be used for BS2 due to reduced penetrance. BS2_Mod is applicable if a single variant positive individual has a negative result for either the IVCT or CHCT diagnostic tests. |

| | |
|---|---|
| **BS3_Moderate** | Well- established *in vitro* or *ex vivo* functional studies or knock-in mouse studies show no damaging effect on protein function.<br>**MHS-*RYR1*:**<br>• Three or more independent *ex vivo* studies all showing **NO** significant increase in release of $Ca^{2+}$ in response to RYR1 agonist (halothane, caffeine, 4-CmC, KCl, voltage). $Ca^{2+}$ release measured by fluorescence. Appropriate controls included. Result must show lack of significant increase in $Ca^{2+}$ release at decreased agonist concentration. |

**SUPPORTING EVIDENCE FOR BENIGN IMPACT**

| | |
|---|---|
| **BP1** | Missense variant in a gene for which primarily truncating variants are known to cause disease.<br>**MHS-*RYR1*:** BP1 is not applicable. MH is caused primarily by missense variants in *RYR1*. |
| **BP2** | Observed in trans with a pathogenic variant for a fully penetrant dominant gene/disorder; or observed in cis with a pathogenic variant in any inheritance pattern.<br>**MHS-*RYR1*:** BP2 is applicable for variants shown to be in cis with a known pathogenic variant. |
| **BP3** | In-frame deletions/insertions in a repetitive region without a known function<br>**MHS-*RYR1*:** BP3 is not applicable. *RYR1* does not have repetitive regions without known function. |
| **BP4** | Multiple lines of computational evidence suggest no impact on gene or gene product (conservation, evolutionary, splicing impact, etc.).<br>**BP4 cannot be used in isolation, at least one other criteria must apply to a variant in order to utilize BP4. If used in isolation it can define a variant as likely benign, it was determined by the VCEP that a variant should not be assessed to be likely benign based solely on computational data.**<br>**MHS-*RYR1*:** REVEL score $\leq$ 0.5 is considered evidence against pathogenicity. |
| **BP5** | Variant found in a case with an alternate molecular basis for disease.<br>**MHS-*RYR1*:** BP5 is not applicable as individuals have been described with MHS and two pathogenic variants in RYR1. |
| **BP6** | Reputable source recently reports variant as benign, but the evidence is not available to the laboratory to perform an independent evaluation.<br>**MHS-*RYR1*:** BP6 has been dropped from the ACMG framework for variant assessment. |
| **BP7** | A synonymous (silent) variant for which splicing prediction algorithms predict no impact to the splice consensus sequence nor the creation of a new splice site AND the nucleotide is not highly conserved.<br>**MHS-*RYR1*:** BP7 applicable as described. |
| **BS3_Supporting** | Well- established *in vitro* or *ex vivo* functional studies or knock-in mouse studies show no damaging effect on protein function.<br>**MHS-*RYR1*:**<br>• **NO** significant increase in $Ca^{2+}$ release in response to RYR1 agonist (halothane, caffeine, 4-CmC, KCl, voltage) in an *in vitro* transfection assay (HEK293, CHO, dyspedic myotubes), $Ca^{2+}$ release measured by fluorescence. Both positive and negative controls included to include variants previously identified as pathogenic and benign. Result must show lack of a significant increase in $Ca^{2+}$ release. Assay must be run in triplicate.<br>• One or two independent *ex vivo* studies all showing **NO** significant increase in release of $Ca^{2+}$ in response to RYR1 agonist (halothane, caffeine, 4-CmC, KCl, voltage). $Ca^{2+}$ release measured by fluorescence. Appropriate controls included. Result must show lack of significant increase in $Ca^{2+}$ release at decreased agonist concentration. |

| | |
|---|---|
| | ◦ *Ex vivo* studies using patient derived samples need to be interpreted with the understanding that unidentified variants may be present. Patient tissues considered useful for BS3 include patient myotubes, microsomal SR preps and lymphoblasts.<br>• **NO** MH reaction in response to RYR1 agonist (halothane, caffeine, 4-CmC, KCl, voltage) in a knock-in mouse model AND **NO** significant increase in sensitivity to RYR1 agonist (halothane, caffeine, 4-CmC, KCl, voltage) in knock-in mouse tissues, $Ca^{2+}$ release measured by fluorescence. Appropriate controls included.<br><br>See attached flow diagram and assay description for assigning PS3/BS3 level of support (Appendix 1). |

**Key: IVCT, in vitro contracture test; CHCT, caffeine halothane contracture test.**

**RULES FOR COMBINING PATHOGENIC CRITERIA**

Bayesian Classification Framework as suggested by Tavtigian et al. 2018 is utilized.

Sum all criteria that are applicable to the variant. Calculate Odds of Pathogenicity using formula below, calculate posterior probability, use posterior probability to determine pathogenicity.

Odds of Pathogenicity = $2.1$ ^ $^{\#\text{Total Supporting}} * 4.3$ ^ $^{\#\text{Total Moderate}} * 18.7$ ^ $^{\#\text{Total Strong}} * 350$ ^ $^{\#\text{Total V Strong}} * 0.4808$ ^ $^{\#\text{Benign Supporting}} * 0.2326$ ^ $^{\#\text{Benign Moderate}} * 0.0535$ ^ $^{\#\text{Benign Strong}}$

Posterior Probability = (Odds Path * 0.1) / (Odds Path - 1) * 0.1+1)

Assignment of Pathogenicity based on Posterior Probability:

| | |
|---|---|
| Posterior Probability < 0.001 | Benign |
| Posterior Probability ≥ 0.001 < 0.1 | Likely Benign |
| Posterior Probability ≥ 0.10 < 0.9 | VUS |
| Posterior Probability ≥ 0.9 to < 0.99 | Likely Pathogenic |
| Posterior Probability ≥ 0.99 | Pathogenic |

Table S2. Review of literature-based data for functional studies showing response of *RYR1* variants to agonist in HEK293 cell transfection assays.

| Variant | Associated Disease State | MH Pathogenicity Assessment without Functional Data | MH Pathogenicity Assessment with Functional Data | Number of Published Assays | Studies Showing $EC_{50}$ Significantly Reduced Compared to WT | Studies Showing $EC_{50}$ NOT Significantly Reduced Compared to WT | Studies Showing No Response to Caffeine and/or Halothane |
|---|---|---|---|---|---|---|---|
| p.(Arg163Cys) | MH | Pathogenic | Pathogenic | 4 | (1-4) | | |
| p.(Gly248Arg) | MH | Pathogenic | Pathogenic | 3 | (1, 4, 5) | | |
| p.(Gly341Arg) | MH | Pathogenic | Pathogenic | 2 | (1, 4) | | |
| p.(Arg401Cys) | MH | Pathogenic | Pathogenic | 2 | (3, 4) | | |
| p.(Arg614Cys) | MH | Pathogenic | Pathogenic | 2 | (1, 4) | | |
| p.(Arg614Leu) | MH | Pathogenic | Pathogenic | 2 | (1, 4) | | |
| p.(Arg2163Cys) | MH | Pathogenic | Pathogenic | 2 | (1, 6) | | |
| p.(Arg2163His) | MH/CCD | Pathogenic | Pathogenic | 2 | (1) | (6) | |
| p.(Val2168Met) | MH | Pathogenic | Pathogenic | 2 | (2, 6) | | |
| p.(Thr2206Met) | MH | Pathogenic | Pathogenic | 2 | (2, 6) | | |
| p.(Arg2336His) | MH | Pathogenic | Pathogenic | 1 | (7) | | |
| p.(Ala2350Thr) | MH | Pathogenic | Pathogenic | 1 | (6) | | |
| p.(Arg2355Trp) | MH | Pathogenic | Pathogenic | 1 | (7) | | |
| p.(Gly2434Arg) | MH | Pathogenic | Pathogenic | 2 | (1, 6) | | |
| p.(Arg2435His) | MH/CCD | Pathogenic | Pathogenic | 2 | (1, 6) | | |
| p.(Arg2454His) | MH | Pathogenic | Pathogenic | 2 | (2, 6) | | |
| p.(Arg2458His) | | Pathogenic | Pathogenic | 2 | (1, 6) | | |
| p.(Arg401His) | MH | Likely Pathogenic | Pathogenic | 1 | (4) | | |
| p.(Tyr522Ser) | MH | Likely Pathogenic | Pathogenic | 2 | (1) | (4) Increase from WT | |
| p.(Arg533Cys) | MH | Likely Pathogenic | Pathogenic | 1 | (3) | | |
| p.(Thr2206Arg) | MH | Likely Pathogenic | Pathogenic | 1 | (2) | | |

| | | | | | | |
|---|---|---|---|---|---|---|
| p.(Gly2375Ala) | MH | Likely Pathogenic | Pathogenic | 1 | (6) | |
| p.(Arg2452Trp) | MH | Likely Pathogenic | Pathogenic | 2 | (5, 8) | |
| p.(Arg2454Cys) | MH | Likely Pathogenic | Pathogenic | 2 | (2, 6) | |
| p.(Arg2458Cys) | MH | Likely Pathogenic | Pathogenic | 2 | (1, 6) | |
| p.(Gly3990Val) | MH | Likely Pathogenic | Pathogenic | 1 | (7) | |
| p.(Val4849Ile) | MH/CCD | Likely Pathogenic | Pathogenic | 2 | (7) | (5) |
| p.(Cys35Arg) | MH | Likely Pathogenic | Likely Pathogenic | 2 | (1) | (4) |
| p.(Arg44Cys) | MH | Likely Pathogenic | Likely Pathogenic | 1 | (3) | |
| p.(Arg163Leu) | MH | Likely Pathogenic | Likely Pathogenic | 1 | (4) | |
| p.(Tyr522Cys) | MH | Likely Pathogenic | Likely Pathogenic | 1 | | (4) |
| p.(Ala2428Thr) | MH | Likely Pathogenic | Likely Pathogenic | 1 | (2) | |
| p.(Arg2508His) | MH/CCD | Likely Pathogenic | Likely Pathogenic | 2 | (9) | (6) |
| p.(Glu3104Lys) | MH | Likely Pathogenic | Likely Pathogenic | 1 | (7) | |
| p.(His4833Tyr) | MH | Likely Pathogenic | Likely Pathogenic | 1 | (3) | |
| p.(Arg328Trp) | MH | VUS | Likely Pathogenic | 1 | (10) | |
| p.(Arg552Trp) | MH | VUS | Likely Pathogenic | 1 | (1) | |
| p.(Ser2345Thr) | MH | VUS | Likely Pathogenic | 1 | (11) | |
| p.(Glu2348del) | MH | VUS | Likely Pathogenic | 1 | (5) | |
| p.(Arg2508Cys) | MH/CCD | VUS | Likely Pathogenic | 3 | (9, 12) | (6) |
| p.(Arg4861His) | MH/CCD | VUS | Likely Pathogenic | 1 | (13) | |
| p.(Thr84Met) | MH | VUS | VUS | 1 | (14) | |
| p.(Thr214Met) | MH DM? | VUS | VUS | 1 | | (5) |
| p.(Arg533His) | MH | VUS | VUS | 1 | (3) | |
| p.(Ser2345Arg) | MH | VUS | VUS | 1 | (11) | |
| p.(Asp3986Glu) | MH | VUS | VUS | 1 | | (7) |
| p.(Ala4894Thr) | MH | VUS | VUS | 1 | (15) | |
| p.(Met4640Ile) | CCD/MHSh | NA | NA | 1 | | (13) |
| p.(Arg2508Gly) | CCD/MHS | NA | NA | 1 | (9) | |
| p.(Tyr4796Cys) | CCD/MHS | NA | NA | 2 | (2, 16) | |
| p.(Pro1787Leu) | NA | Benign_BA1 | Benign_BA1 | 2 | | (2, 17) |
| p.(Gly2060Cys) | NA | Benign_BA1 | Benign_BA1 | 1 | | (17) |
| p.(Arg2508Lys) | NA | NA | NA | 1 | (9) | |

| | | | | | | | |
|---|---|---|---|---|---|---|---|
| p.(Ala4894Ser) | NA | NA | NA | 1 | (15) | | |
| p.(Ala4894Gly) | NA | NA | NA | 1 | | (15)<br>$EC_{50}$<br>INCREASED | |
| p.(Leu4647Pro) | CCD | NA | NA | 1 | | | (17) |
| p.(Phe4857Ser) | CCD | NA | NA | 1 | | | (13) |
| p.(Ala4894Pro) | Myopathy | NA | NA | 1 | | | (15) |
| p.(Phe4808Leu) | CCD | NA | NA | 1 | | | (17) |
| p.(Asp4918Asn) | CCD | NA | NA | 2 | | | (13, 17) |
| p.(Arg4893Gln) | CCD | NA | NA | 1 | | | (17) |
| p.(Ile403Met) | CCD | NA | NA | 1 | (1) | | |
| p.(Lys3367Arg) | CCD | NA | NA | 1 | | (11) | |

Key: MH, malignant hyperthermia; MHS, malignant hyperthermia susceptibility based on *in vitro* contracture test (IVCT) or caffeine halothane contracture test (CHCT); MHSh, malignant hyperthermia susceptibility with sensitivity limited to halothane; CCD, central core disease; NA, not applicable.

Table S3. ACMG criteria and pathogenicity assessment of 44 variants on the European Malignant Hyperthermia Group list of diagnostic mutations.

| Genomic Coordinate (GRCh37) | cDNA NM_000540.2 | Protein | ACMG/AMP Criteria | REVEL | SIFT | Pathogenicity | Posterior Probability | References |
|---|---|---|---|---|---|---|---|---|
| chr19:g.38931442T>C | c.103T>C | p.(Cys35Arg) | PS4_Mod, PM1, PP1_St, PP3_Mod | 0.947 | D | Pathogenic | 0.994 | (1, 2, 4, 18-24) |
| chr19:g.38931469C>T | c.130C>T | p.(Arg44Cys) | PS3_Mod, PS4_Mod, PM1, PP3_Mod | 0.951 | D | Likely Pathogenic | 0.975 | (3, 19, 25, 26) |
| chr19:g.38934851C>T | c.487C>T | p.(Arg163Cys)[a,b] | PS3, PS4, PM1, PM5, PP1_St, PP3_Mod | 0.959 | D | Pathogenic | 1.000 | (1-4, 22-24, 27-59) |
| chr19:g.38934852G>T | c.488G>T | p.(Arg163Leu) | PS3_Mod, PS4_Mod, PM1, PP3_Mod | 0.882 | D | Likely Pathogenic | 0.975 | (2, 4, 22, 56, 60) |
| chr19:g.38937350G>C | c.742G>C | p.(Gly248Arg)[a,b] | PS1, PS3_Mod, PS4_Mod, PM1, PP1_Mod, PP3_Mod | 0.883 | D | Pathogenic | 1.000 | (4, 5, 22, 24, 29, 34, 48, 61, 62) |
| chr19:g.38937350G>A | c.742G>A | p.(Gly248Arg) | PS3_Mod, PS4, PM1, PP1, PP3_Mod | 0.889 | D | Pathogenic | 0.997 | (1, 2, 4, 22, 24, 29, 31, 43, 50, 51, 62-64) |
| chr19:g.38939313C>T | c.982C>T | p.(Arg328Trp) | PS3_Mod, PS4_Supp, PM1, PP1 | 0.76 | D | Likely Pathogenic | 0.900 | (2, 10) |
| chr19:g.38939352G>A | c.1021G>A | p.(Gly341Arg)[a,b] | PS3_Mod, PS4, PM1, PP1_St, PP3_Mod | 0.864 | D | Pathogenic | 1.000 | (2, 4, 18, 19, 22, 23, 26, 27, 30-34, 36, 41, 43, 46, 47, 50, 52, 53, 65-76) |
| chr19:g.38939352G>C | c.1021G>C | p.(Gly341Arg)[a,b] | PS1, PS3_Mod, PS4_Mod, PM1, PP1_St, PP3_Mod | 0.876 | D | Pathogenic | 1.000 | (1, 2, 4, 19, 22, 24, 26, 30, 39, 45, 47, 74) |

| | | | | | | | | |
|---|---|---|---|---|---|---|---|---|
| chr19:g.38942482C>T | c.1201C>T | p.(Arg401Cys)[a,b] | PS3_Mod, PS4_Mod, PM1, PM5, PP1_Mod, PP3_Mod | 0.886 | D | Pathogenic | 0.999 | (2, 3, 22, 26, 34, 44, 46, 52, 77, 78) |
| chr19:g.38945999A>C | c.1565A>C | p.(Tyr522Ser) | PS3, PS4_Supp, PM1, PP1 PP3_Mod | 0.942 | D | Pathogenic | 0.994 | (1, 2, 4, 22, 24, 42, 43, 79-85) |
| chr19:g.38946103G>A | c.1589G>A | p.(Arg530His) | PS4_Supp, PM1, PP3_Mod | 0.93 | D | VUS | 0.812 | (22, 86, 87) |
| chr19:g.38946111C>T | c.1597C>T | p.(Arg533Cys) | PS3_Mod, PS4_Supp, PM1, PP1_St, PP3_Mod | 0.918 | D | Pathogenic | 0.997 | (3, 19, 25) |
| chr19:g.38946112G>A | c.1598G>A | p.(Arg533His) | PS3_Mod, PM1 | 0.824 | D | VUS | 0.675 | (3, 22, 39) |
| chr19:g.38946168C>T | c.1654C>T | p.(Arg552Trp) | PS3_Mod, PS4_Mod, PM1, PP1 | 0.831 | D | Likely Pathogenic | 0.949 | (1, 2, 22-24, 43, 60, 88, 89) |
| chr19:g.38948185C>T | c.1840C>T | p.(Arg614Cys)[a,b] | PS3_Mod, PS4, PM5, PP1_St, PP3_Mod | 0.927 | D | Pathogenic | 1.000 | (1, 2, 4, 18, 19, 22-24, 26, 27, 29-31, 33, 34, 36, 37, 41-43, 45-50, 52-54, 57, 60, 67, 68, 74-77, 87, 90-114) |
| chr19:g.38948186G>T | c.1841G>T | p.(Arg614Leu)[a,b] | PS3_Mod, PS4, PP1_St, PP3_Mod | 0.931 | D | Pathogenic | 0.999 | (1, 2, 4, 18, 19, 22-24, 26, 33, 43, 61, 75, 90, 114) |
| chr19:g.38985204C>T | c.6487C>T | p.(Arg2163Cys)[a,b] | PS3_Mod, PS4_Mod, PM1, PM5, PP1_St, PP3_Mod | 0.956 | D | Pathogenic | 1.000 | (1, 2, 6, 18, 22-24, 32, 33, 43, 44, 47-49, 57, 90, 109, 110) |

| | | | | | | | | |
|---|---|---|---|---|---|---|---|---|
| chr19:g.38985205G>A | c.6488G>A | p.(Arg2163His)[a,b] | PS3_Mod, PS4, PM1, PP1_St, PP3_Mod | 0.933 | D | Pathogenic | 1.000 | (1, 2, 6, 22-25, 31-33, 36, 39, 42-44, 49, 50, 53, 109, 115) |
| chr19:g.38985219G>A | c.6502G>A | p.(Val2168Met)[a,b] | PS3_Mod, PS4, PM1, PP1_St, PP3_Mod | 0.896 | D | Pathogenic | 1.000 | (2, 6, 18, 19, 22, 23, 25, 26, 29-31, 46-48, 50, 53, 68, 77, 87, 89-91, 107, 108, 116-118) |
| chr19:g.38986923C>T | c.6617C>T | p.(Thr2206Met)[a,b] | PS3_Mod, PS4, PM1, PM5, PP1_St, PP3_Mod | 0.95 | D | Pathogenic | 1.000 | (2, 6, 22, 23, 26, 29-32, 34, 44, 46, 48-50, 67, 68, 74, 75, 77, 90, 97, 117, 119-121) |
| chr19:g.38986923C>G | c.6617C>G | p.(Thr2206Arg) | PS3_Mod, PS4_Mod, PM1, PP1_Mod, PP3_Mod | 0.968 | D | Pathogenic | 0.994 | (2, 22, 23, 30, 46, 49, 52, 89, 90, 117, 119) |
| chr19:g.38989863G>A | c.7007G>A | p.(Arg2336His)[a,b] | PS3_Mod, PS4, PM1, PP1_St, PP3_Mod | 0.903 | D | Pathogenic | 0.999 | (7, 22, 26, 31, 36, 49, 50, 53, 68, 75, 86, 122) |
| chr19:g.38990289_ 38990291del | c.7042_7044del | p.(Glu2348del) | PS3_Mod, PS4_Mod, PM1 | NA | | Likely Pathogenic | 0.900 | (5, 22, 31, 109, 115, 123) |
| chr19:g.38990295G>A | c.7048G>A | p.(Ala2350Thr)[a,b] | PS3_Mod, PS4, PM1, PP1_Mod, PP3_Mod | 0.952 | D | Pathogenic | 0.999 | (2, 6, 22, 26, 32, 46-48, 50, 67, 75, 95, 124-126) |

| | | | | | | | | |
|---|---|---|---|---|---|---|---|---|
| chr19:g.38990310C>T | c.7063C>T | p.(Arg2355Trp)[a,b] | PS3_Mod, PS4, PM1, PP1_St, PP3_Mod | 0.861 | D | Pathogenic | 1.000 | (7, 22, 31, 36, 50, 125, 127-129) |
| chr19:g.38990371G>C | c.7124G>C | p.(Gly2375Ala) | PS3_Mod, PS4_Mod, PM1, PP1_Mod, PP3_Mod | 0.905 | D | Pathogenic | 0.994 | (6, 26, 46, 74, 75, 125, 130-132) |
| chr19:g.38990615G>A | c.7282G>A | p.(Ala2428Thr) | PS3_Mod, PS4_Mod, PM1, PP3_Mod | 0.85 | D | Likely Pathogenic | 0.975 | (2, 22, 30, 46, 76) |
| chr19:g.38990633G>A | c.7300G>A | p.(Gly2434Arg)[a,b] | PS3, PS4, PM1, PP1_St, PP3_Mod | 0.965 | D | Pathogenic | 1.000 | (1, 2, 6, 19, 22, 24, 26, 27, 29-36, 43, 44, 46, 48-50, 52, 53, 64, 67, 68, 74, 75, 77, 90, 91, 97, 99, 104, 107, 108, 115, 133-138) |
| chr19:g.38990637G>A | c.7304G>A | p.(Arg2435His)[a,b] | PS2/PM6_Mod, PS3_Mod, PS4, PM1, PP1_St, PP3_Mod | 0.944 | D | Pathogenic | 1.000 | (1, 2, 6, 18, 19, 22, 24, 30, 35, 39, 42, 43, 46, 49, 50, 62, 67, 68, 97, 99, 115, 129, 136, 139-141) |
| chr19:g.38991276C>T | c.7354C>T | p.(Arg2452Trp)[a] | PS3_Mod, PS4_Mod, PM1, PP1_St | 0.828 | D | Pathogenic | 0.994 | (5, 8, 22, 26, 30, 46, 67, 95, 142, 143) |
| chr19:g.38991282C>T | c.7360C>T | p.(Arg2454Cys) | PS3_Mod, PS4_Mod, PM1, PM5, PP1, PP3_Mod | 0.913 | D | Pathogenic | 0.997 | (2, 6, 22, 26, 45, 75, 90, 144) |

| | | | | | | | | |
|---|---|---|---|---|---|---|---|---|
| chr19:g.38991283G>A | c.7361G>A | p.(Arg2454His)[a,b] | PS3_Mod, PS4, PM1, PP1_St, PP3_Mod | 0.923 | D | Pathogenic | 1.000 | (2, 6, 19, 22, 25, 26, 29-31, 33, 36, 46, 48, 53, 67, 87, 90, 97, 115, 142) |
| chr19:g.38991294C>T | c.7372C>T | p.(Arg2458Cys) | PS3_Mod, PS4_Mod, PM1, PM5, PP3_Mod | 0.922 | D | Pathogenic | 0.994 | (1, 2, 6, 19, 22, 24, 26, 32, 33, 43, 44, 49, 91, 107, 108, 145) |
| chr19:g.38991295G>A | c.7373G>A | p.(Arg2458His)[a,b] | PS3_Mod, PS4, PM1, PP1_St, PP3_Mod | 0.959 | D | Pathogenic | 1.000 | (1, 2, 6, 22, 24, 34, 39, 43, 47, 50, 62, 145-148) |
| chr19:g.38991538C>T | c.7522C>T | p.(Arg2508Cys) | PS3_Mod, PS4_Mod, PP3_Mod | 0.861 | D | Likely Pathogenic | 0.900 | (6, 9, 12, 22, 39, 49, 62, 141, 149) |
| chr19:g.38991539G>A | c.7523G>A | p.(Arg2508His) | PS2/PM6_Supp, PS3_Mod, PS4, PP3_Mod | 0.898 | D | Likely Pathogenic | 0.988 | (6, 9, 22, 31, 36, 39, 49, 67, 68, 74, 141) |
| chr19:g.39002961G>A | c.9310G>A | p.(Glu3104Lys) | PS3_Mod, PS4_Mod, PP1_Mod, PP3_Mod | 0.868 | D | Likely Pathogenic | 0.975 | (7, 22, 50) |
| chr19:g.39034472G>T | c.11969G>T | p.(Gly3990Val)[a,b] | PS3_Mod, PS4, PP1_St | 0.75 | D | Pathogenic | 0.995 | (7, 22, 50, 52) |
| chr19:g.39070734C>T | c.14477C>T | p.(Thr4826Ile)[a,b] | PS3, PS4, PM1_Sup, PP1_St, PP3_Mod | 0.977 | D | Pathogenic | 1.000 | (2, 21, 22, 34, 47, 50, 51, 57, 67, 76, 109, 110, 115, 150, 151) |
| chr19:g.39070754C>T | c.14497C>T | p.(His4833Tyr) | PS3_Mod, PS4_Mod, | 0.975 | D | Likely Pathogenic | 0.988 | (3, 51, 152, 153) |

| | | | | | | | | |
|---|---|---|---|---|---|---|---|---|
| | | | PM1_Sup, PP1_Mod, PP3_Mod | | | | | |
| chr19:g.39071010C>G | c.14512C>G | p.(Leu4838Val) | PS3_Supp, PS4_Mod, PM1_Sup, PP3_Mod | 0.882 | D | Likely Pathogenic | 0.900 | (2, 22, 39, 62, 73, 86, 115, 154) |
| chr19:g.39071043G>A | c.14545G>A | p.(Val4849Ile)[a,b] | PS3_Mod, PS4, PM1_Supp, PP1_St | 0.817 | D | Pathogenic | 0.997 | (2, 7, 13, 22, 26, 31, 32, 35, 50, 52, 53, 61, 67, 141, 155) |
| chr19:g.39071080G>A | c.14582G>A | p.(Arg4861His) | PS3_Mod, PS4_Sup, PM1_Sup, PP3_Mod | 0.912 | D | Likely Pathogenic | 0.900 | (13, 22, 31, 67, 97, 141) |

Key: ACMG/AMP codes are the codes from Richards et al (2015), REVEL score is a metapredictor score.(156)

[a]Variants assessed to be pathogenic that were used in likelihood ratio calculations for PP3/BP4 (computational) and for [b]PM1 (hotspot) weighting. For weighting of the PM1 criterion, only variants that reached an assessment of pathogenic without the use of PM1 were considered (21 variants). For weighting of PP3/BP4 only variants that reached an assessment of pathogenic without the use of PP3 were considered (22 variants). NA, not applicable.

Table S4. *RYR1* variants in gnomAD determined to be benign based on popmax frequency, BA1 assigned for popmax freq ≥0.0038. These variants were used for likelihood calculations for PM1 (hotspot) and PP3/BP4 (computational) weighting.

| Genomic Coordinate (GRCh37) | cDNA NM_000540.2 | Protein | REVEL | SIFT | PopMax* MAF | PopMax* | gnomAD MAF |
|---|---|---|---|---|---|---|---|
| chr19:g.38954162G>A | c.2677G>A | p.(Gly893Ser) | 0.837 | D | 0.0070 | AFR | 0.0007 |
| chr19:g.38958397G>A | c.3326G>A | p.(Arg1109Lys) | 0.088 | T | 0.0189 | AFR | 0.0019 |
| chr19:g.38964275A>G | c.4024A>G | p.(Ser1342Gly) | 0.293 | T | 0.1566 | AFR | 0.0159 |
| chr19:g.38964306C>G | c.4055C>G | p.(Ala1352Gly) | 0.274 | T | 0.0222 | AFR | 0.0023 |
| chr19:g.38965975A>G | c.4178A>G | p.(Lys1393Arg) | 0.555 | T | 0.0046 | NFE | 0.0027 |
| chr19:g.38974116C>T | c.4894C>T | p.(Pro1632Ser) | 0.821 | D | 0.0201 | AFR | 0.0020 |
| chr19:g.38976294C>T | c.4999C>T | p.(Arg1667Cys) | 0.339 | D | 0.0051 | EAS | 0.0020 |
| chr19:g.38976612C>T | c.5317C>T | p.(Pro1773Ser) | 0.46 | D | 0.0147 | EAS | 0.0011 |
| chr19:g.38976655C>T | c.5360C>T | p.(Pro1787Leu) | 0.055 | T | 0.0413 | SAS | 0.0188 |
| chr19:g.38979903G>C | c.5634G>C | p.(Glu1878Asp) | 0.061 | T | 0.0150 | AFR | 0.0015 |
| chr19:g.38983180G>T | c.6178G>T | p.(Gly2060Cys) | 0.172 | T | 0.1562 | SAS | 0.0660 |
| chr19:g.38995998C>G | c.8360C>G | p.(Thr2787Ser) | 0.466 | D | 0.0299 | AFR | 0.0035 |
| chr19:g.38998362G>A | c.8827G>A | p.(Asp2943Asn) | 0.725 | D | 0.0110 | AFR | 0.0010 |
| chr19:g.39002893T>C | c.9242T>C | p.(Met3081Thr) | 0.738 | D | 0.0071 | AFR | 0.0007 |
| chr19:g.39003004C>T | c.9353C>T | p.(Ala3118Val) | 0.213 | D | 0.0063 | AFR | 0.0006 |
| chr19:g.39018347G>C | c.10747G>C | p.(Glu3583Gln) | 0.32 | T | 0.0241 | SAS | 0.0147 |
| chr19:g.39019242C>G | c.10941C>G | p.(His3647Gln) | 0.51 | T | 0.0115 | AFR | 0.0010 |
| chr19:g.39025366C>G | c.11266C>G | p.(Gln3756Glu) | 0.326 | T | 0.1085 | AMR | 0.0331 |
| chr19:g.39034444C>T | c.11941C>T | p.(His3981Tyr) | 0.494 | T | 0.0137 | AFR | 0.0014 |
| chr19:g.39055821G>A | c.12847G>A | p.( Glu4283Lys) | 0.188 | T | 0.0183 | AMR | 0.0006 |
| chr19:g.39055843C>T | c.12869C>T | p.(Ala4290Val) | 0.27 | D | 0.0078 | AFR | 0.0027 |
| chr19:g.39055854A>G | c.12880A>G | p.(Thr4294Ala) | 0.251 | T | 0.0155 | EAS | 0.0010 |
| chr19:g.39055855C>T | c.12881C>T | p.(Thr4294Met) | 0.528 | T | 0.0139 | AFR | 0.0047 |
| chr19:g.39055930G>A | c.12956G>A | p.(Arg4319Gln) | 0.323 | T | 0.0078 | AFR | 0.0028 |
| chr19:g.39057615C>T | c.13502C>T | p.(Pro4501Leu) | 0.418 | D | 0.0181 | AFR | 0.0025 |
| chr19:g.39057626G>C | c.13513G>C | p.(Asp4505His) | 0.663 | D | 0.0054 | NFE | 0.0033 |
| chr19:g.39062759C>A | c.13847C>A | p.(Ala4616Asp) | 0.225 | T | 0.0047 | AFR | 0.0004 |

*PopMax refers to the outbred population with the highest minor allele frequency in gnomAD. MAF, minor allele frequency. T, tolerated; D, damaging.

Table S5. Assessment of forty *RYR1* variants using revised ACMG/AMP guidelines. Variants selected from ClinVar, all variants were assessed for malignant hyperthermia by at least one submitter.

| Genomic Position 37 | cDNA NM_000540.2 | Protein | Prior ClinVar Pathogenicity | ClinVar Accession | ACMG/AMP Criteria | REVEL Score | VCEP Pathogenicity | Posterior Probability | References |
|---|---|---|---|---|---|---|---|---|---|
| Chr19:g.38931491C>A | c.152C>A | p.(Thr51Asn) | LB | VCV000133099 | PM1 | 0.769 | VUS | 0.325 | (34) |
| Chr19:g.38945887A>G | c.1453A>G | p.(Met485Val) | B/LB/VUS | VCV000133076 | PM1 | 0.551 | VUS | 0.325 | (157-159) |
| Chr19:g.38946111C>A | c.1597C>A | p.(Arg533Ser) | LP | VCV000291315 | PM1, PP3_Mod | 0.86 | VUS | 0.675 | |
| Chr19:g.38948887G>A | c.2122G>A | p.(Asp708Asn) | B/VUS | VCV000159840 | NA | 0.726 | VUS | 0.100 | (35, 159) |
| Chr19:g.38955289G>A | c.2797G>A | p.(Ala933Thr) | LB/B | VCV000161363 | BA1 | 0.926 | VUS | <0.001 | (160) |
| Chr19:g.38956856G>A | c.2996G>A | p.(Arg999His) | LB | VCV000133122 | NA | 0.817 | VUS | 0.100 | (161) |
| Chr19:g.38964275A>G | c.4024A>G | p.(Ser1342Gly) | B | VCV000093265 | BA1 | 0.293 | Benign | <0.001 | (22, 60, 77, 86, 112, 160, 162, 163) |
| Chr19:g.38968456A>G | c.4400A>G | p.(Lys1467Arg) | VUS | VCV000161371 | BP4 | 0.371 | VUS | 0.051 | (19) |
| Chr19:g.38973933A>G | c.4711A>G | p.(Ile1571Val) | B/VUS | VCV000159851 | BS1 | 0.56 | Likely Benign | 0.006 | (19, 60, 67, 77, 155, 159, 164) |
| Chr19:g.38976331G>A | c.5036G>A | p.(Arg1679His) | LB/B | VCV000161364 | PP3_Mod, BS1 | 0.918 | Likely Benign | 0.025 | (35, 61, 165, 166) |
| Chr19:g.38976478C>T | c.5183C>T | p.(Ser1728Phe) | P | VCV000133144 | PS4, PP1_Mod, BP4 | 0.477 | VUS | 0.812 | (22, 32, 50, 52, 163) |
| Chr19:g.38976612C>T | c.5317C>T | p.(Pro1773Ser) | B/VUS | VCV000224382 | BA1 | 0.46 | Benign | 0.051 | (39) |
| Chr19:g.38976655C>T | c.5360C>T | p.(Pro1787Leu) | B | VCV000133149 | BA1 | 0.055 | Benign | <0.001 | (2, 22, 31, 60, 63, 77, 86, 87, 163, 167) |
| Chr19:g.38983180G>T | c.6178G>T | p.(Gly2060Cys) | B | VCV000093279 | BA1 | 0.172 | Benign | <0.001 | (19, 22, 26, 31, 60, 63, 77, 86, 87, 155, 163, 166, 167) |

| | | | | | | | | | |
|---|---|---|---|---|---|---|---|---|---|
| Chr19:g.38985195G>A | c.6478G>A | p.(Gly2160Ser) | VUS | | PM1 | 0.635 | VUS | 0.325 | (22, 160) |
| Chr19:g.38987056G>A | c.6671G>A | p.(Arg2224His) | LP | VCV000478260 | PM1 | 0.605 | VUS | 0.325 | |
| Chr19:g.38990320T>A | c.7073T>A | p.(Ile2358Asn) | LP | VCV000803553 | PM1, PP3_Mod | 0.867 | VUS | 0.675 | |
| Chr19:g.38990370G>A | c.7123G>A | p.(Gly2375Arg) | LP | VCV000590582 | PS4_Sup, PM1, PM5, PP3_Mod | 0.9 | Likely Pathogenic | 0.949 | (22, 74) |
| Chr19:g.38991307C>T | c.7385C>T | p.(Pro2462Leu) | LB | VCV000329061 | PP3_Mod | 0.883 | VUS | 0.325 | |
| Chr19:g.39008071T>C | c.9758T>C | p.(Ile3253Thr) | B/VUS | VCV000159865 | NA | 0.845 | VUS | 0.100 | (31, 60) |
| Chr19:g.39009877C>T | c.10042C>T | p.(Arg3348Cys) | VUS | VCV000329095 | PS4_Sup | 0.784 | VUS | 0.188 | (22, 146) |
| Chr19:g.39009932G>A | c.10097G>A | p.(Arg3366His) | B/VUS | VCV000132990 | BS1 | 0.68 | Likely Benign | 0.006 | (60, 67, 77, 155, 164) |
| Chr19:g.39016072C>T | c.10556C>T | p.(Pro3519Leu) | LP | VCV000803555 | NA | 0.809 | VUS | 0.100 | |
| Chr19:g.39018347G>C | c.10747G>C | p.(Glu3583Gln) | B | VCV000132999 | BA1 | 0.32 | Benign | <0.001 | (22, 31, 60, 155, 163) |
| Chr19:g.39025366C>G | c.11266C>G | p.(Gln3756Glu) | B | VCV000133011 | BA1 | 0.326 | Benign | <0.001 | (22, 32, 39, 60, 64, 77, 87, 150, 154, 163, 168) |
| Chr19:g.39025414C>T | c.11314C>T | p.(Arg3772Trp) | P | VCV000478159 | PS4_Sup, PM5, PP3_Mod | 0.939 | VUS | 0.812 | (86, 112, 169) |
| Chr19:g.39025415G>A | c.11315G>A | p.(Arg3772Gln) | P | VCV000133012 | PS4, PP1_Strong, PP3_Mod | 0.888 | Pathogenic | 0.994 | (22, 159, 170) |
| Chr19:g.39034191A>G | c.11798A>G | p.(Tyr3933Cys) | B/LB/VUS | VCV000133021 | PP3_Mod, BS1 | 0.983 | Likely Benign | 0.025 | (19, 31, 34, 60, 67, 77, 155, 159, 164) |
| Chr19:g.39034450C>T | c.11947C>T | p.(Arg3983Cys) | LP | VCV000650932 | PS2_PM6_Mod, PS4_Sup, PP3_Mod, BS3_Sup | 0.931 | VUS | 0.675 | (121, 171) |
| Chr19:g.39034461C>G | c.11958C>G | p.(Asp3986Glu) | P | VCV000133026 | PS4, PP1, BS3_Sup | 0.763 | VUS | 0.675 | (7, 22, 31, 50, 60, 163) |
| Chr19:g.39052002G>A | c.12532G>A | p.(Gly4178Ser) | LP | VCV000374083 | PS4_Sup, PP3_Mod | 0.979 | VUS | 0.500 | (26, 77) |

| | | | | | | | | | |
|---|---|---|---|---|---|---|---|---|---|
| Chr19:g.39055855C>T | c.12881C>T | p.(Thr4294Met) | VUS | VCV000159834 | BA1 | 0.528 | Benign | <0.001 | (160) |
| Chr19:g.39055858C>T | c.12884C>T | p.(Ala4295Val) | VUS | VCV000133043 | BS1, BP4 | 0.152 | Likely Benign | 0.003 | (22, 60, 62, 139, 159) |
| Chr19:g.39057618A>G | c.13505A>G | p.(Glu4502Gly) | B/LB/VUS | VCV000161366 | NA | 0.532 | VUS | 0.100 | |
| Chr19:g.39057626G>C | c.13513G>C | p.(Asp4505His) | B/LB/VUS | VCV000093252 | BA1 | 0.663 | Benign | <0.001 | (22, 26, 77, 159) |
| Chr19:g.39061260G>A | c.13673G>A | p.(Arg4558Gln) | LB | VCV000065984 | PP3_Mod | 0.957 | VUS | 0.325 | |
| Chr19:g.39061289C>G | c.13702C>G | p.(Leu4568Val) | LP | VCV000803556 | NA | 0.775 | VUS | 0.100 | |
| Chr19:g.39062830A>G | c.13918A>G | p.(Met4640Val) | LP | VCV000803557 | PS4_Sup, PM1_Sup, PP3_Mod | 0.911 | VUS | 0.675 | (13) |
| Chr19:g.39063944C>T | c.14126C>T | p.(Thr4709Met) | LP/VUS | VCV000065996 | PM1_Sup, PP3_Mod, BS3_Sup | 0.901 | VUS | 0.325 | (159) |
| Chr19:g.39076780C>T | c.14918C>T | p.(Pro4973Leu) | LP | VCV000133098 | PS4_Mod, PM1_Sup, PP3_Mod | 0.898 | VUS | 0.812 | (2, 19, 22, 31, 44, 45, 115, 163, 172) |

Key: B benign, LB, likely benign, VUS variant of uncertain significance, LP, likely pathogenic, P, pathogenic. ACMG/AMP codes are the codes from Richards et al (2015), REVEL score is a metapredictor score.(156)

**Supplemental Information**

**Supplemental Methods**

**Data collection methods:** The HGMD (https://portal.biobase-international.com/hgmd/pro/start.php), ClinVar (https://www.ncbi.nlm.nih.gov/clinvar/), GoogleScholar (https://scholar.google.com) and MasterMind (https://mastermind.genomenon.com) were utilized to identify relevant articles.

**Case Information:** Results from either the Caffeine Halothane Contracture Test (CHCT) or the *In Vitro* Contracture Test (IVCT) were considered valid diagnostic tests for determination of MH status. Both tests rely on isolating muscle fibers from affected individuals and determining the strength of contraction when the fiber is exposed to caffeine and halothane. Individuals are determined to be MH-susceptible (MHS), or MH-negative (MHN). Only individuals who were diagnosed as MHS were considered to have a positive test for the purpose of variant assessment.

**Explanation for criteria dropped from RYR1 specific guidelines:** Because the *RYR1/*MH pathogenic variant spectrum consists almost entirely of missense variants, with just a handful of small, in-frame deletions and no loss of function variants the following criteria were dropped; PVS1 (putative loss of function variants), PM4 (protein length change), and BP1 (missense variant in gene where predominantly loss of function variants cause disease). Because MH typically follows an autosomal dominant inheritance pattern, we dropped PM3 (in *trans* to pathogenic variant for recessive disorder). Because *RYR1* missense variants are not uncommon in gnomAD, we dropped PP2 (low rate of benign missense variants in gene). Because multiple loci contribute to *RYR1*, we dropped PP4 (patient's phenotype specific for disease with single genetic etiology). It has been reported that up to 12% of family members that test positive by IVCT/CHCT are negative for the familial *RYR1*/MH variant. Therefore, we dropped BS4 (lack of segregation). While it has been suggested that the CHCT/IVCT diagnostic test has a high false positive rate, it is possible that other MH pathogenic variants exist in these families that account for this finding (Miller et al. 2018) and it was not felt to be a reliable criterion for this disorder. Because *RYR1* does not have regions of sequence repeats, we dropped BP3 (variant in repetitive region without known function). There have been reports of individuals with MHS who have been found to have two pathogenic *RYR1* variants (Kraeva et al. 2011) and at least one family has been identified with pathogenic variants in both *RYR1* and *CACNA1S*.(Monnier et al. 2002) Based on these reports, we dropped the BP5 criterion (alternate molecular basis for disease).

**How to evaluate evidence strength for Bayesian pathogenicity criteria using likelihood ratios.**

The general form of the Bayes equation is:

$$P(A|B) = P(B|A)*P(A)/P(B)$$

In pathogenicity assessments:

A is the pathogenicity

B is the evidence

So, P(Path|Evid) is “The probability of pathogenicity given the applied evidence”, which is what we are setting out to determine. The vertical bar (pipe) means ‘given’. This is also called the posterior probability, that is the pathogenicity of the variant after applying the evidence.

To do that we make a calculation, which is based on three factors:

P(Evid|Path) is “The probability of the evidence, given a pathogenic variant”
P(Path) is “The probability of a pathogenic variant”
P(Evid) is “The probability of observing the evidence”

To make things even easier, this equation can be rearranged and simplified using some unfamiliar, but very handy terms.

Prior Probability – the likelihood of observing something before a piece of evidence is observed.
Conditional Probability – this is P(Evid|Path) – the likelihood of observing the evidence if the variant is pathogenic. This can be expressed in the form of the Odds of Pathogenicity for a given piece of evidence, which we call OddsP

Posterior Probability – this is P(Path|Evid)

Prob(Path|Evid) = (OddsP*Prior)/((OddsP-1)*Prior+1)

From Tavtigian et al, the ‘OddsP’ is equal to the product of the Odds of pathogenicity of all of the criteria that one would use in the assessment.
In the current Bayesian formulation of the ACMG/AMP criteria, the prior probability is 0.1.

Richards et al implicitly set relative OddsP for each criterion, using the categories of ‘Very Strong’, ‘Strong’, ‘Moderate’, and ‘Supporting’. Tavtigian et al transformed those into OddsP of 350:1, 18.7:1, 4.3:1, and 2.08:1

In this system, the OddsP of each criterion is multiplied together to determine the overall OddsP. Here are some examples of how to estimate the strength of these OddsP for each of several criteria.

**In silico data analysis**
Let us say we are trying to evaluate the strength of evidence for an *in silico* predictor for variants in RYR1 because we want to know how to use that evidence. For simplicity I will illustrate this using SIFT. The simplest way to do this is to just use the categorical output of SIFT as recommended by the programmers – either ‘Damaging’ or ‘Tolerated’. To evaluate the SIFT

predictor strength for PP3/BP4, we look at a set of variants we are confident are pathogenic and another set that we are confident are benign. For 22 known pathogenic RYR1 variants, ~100% of the time (22 variants) the *in silico* predictor reads out damaging, and ~0% of the time (0 variants) it reads out 'Tolerated'. For known benign variants, 41% of the time it reads out damaging (11 variants), and 59% of the time (16 variants) it reads out tolerated.

SIFT for the RYR1 sample set of 29 pathogenic variants and 27 benign variants.

| | Path | Benign |
|---|---|---|
| Damaging | 22 | 11 |
| Tolerated | 1* | 16 |

The likelihood ratio for a Damaging readout (LR+) or a Tolerated readout (LR-) for a variant of unknown pathogenicity requires use of the Sensitivity and Specificity of the in silico predictor.
*We need a "1" for this position to allow for the calculations to work.

Sensitivity is:
True Positives/(True Positives + False Negatives) = 22/(22+1) = 0.957

Specificity is:
True Negatives/(True Negatives + False Positives) = 16/(16+11) = 0.593

LR+ = Sensitivity/(1-Specificity) =
{(TP/(TP+FN)}/[{(1-{TN/(TN+FP)}] =
{(22/(22+1)}/[{(1-{16/(16+11)}] = 2.35, or 2.35:1

Note that the online LR calculators allow you to derive this directly from the 2 x 2 table.

So, the OddsP for a SIFT readout of 'Damaging' is between Supporting (2.08:1) and Moderate (4.3:1) and should be used as Supporting. Note that an LR+ of 1.0 would be no data for or against pathogenicity – the odds are 1:1. When you use the online LR calculators, they also give you the 95% confidence interval of the LR, which in this case is 1.50-3.73. It is a bit worrisome that the lower bound of the CI is below the specified LR for supporting evidence (2.08:1), suggesting that this limited dataset does not provide robust support for implementing this as supportive evidence. There is debate about this as to whether one should use point estimates for such data or the conservative bound of a 95% CI, but that is beyond the scope of this guide.

The likelihood ratio for a Tolerant readout (LR-) for a variant of unknown pathogenicity would be:

LR- = (1-Sensitivity)/Specificity = [1-{TP/(TP+FN)}]/{TN/(TN+FP)} = [1-{22/(22+1)}]/{16/(16+11)} = 0.07

So, the OddsP for a SIFT readout of 'Tolerant' is~ 0.07, or about 1:14 *for* pathogenicity, or 14:1 *against* pathogenicity, or 14:1 *for* benign. This would lie between Moderate (4.3:1) and Strong (18.7:1). Note that Richards et al did not have an evidence level of Moderate for Benign evidence. So, in that framework one would have to use this as benign supporting, or perhaps two benign supporting pieces of evidence. In Tavtigian et al, Formula 5 could be amended from the published form:

$$OP = O_{VSt}^{\left(\frac{N_{PSu}}{8}+\frac{N_{PM}}{4}+\frac{N_{PSt}}{2}+\frac{N_{PVSt}}{1}-\frac{N_{BSu}}{8}-\frac{N_{BSt}}{2}\right)}$$

To:

$$OP = O_{VSt}^{\left(\frac{N_{PSu}}{8}+\frac{N_{PM}}{4}+\frac{N_{PSt}}{2}+\frac{N_{PVSt}}{1}-\frac{N_{BSu}}{8}-\frac{N_{BMo}}{4}-\frac{N_{BSt}}{2}\right)}$$

By adding the term highlighted in yellow, which counts Moderate Benign evidence, this evidence strength can then be calculated into the overall pathogenicity .

One can also consider a trichotomization* of *in silico* data, rather than dichotomization. In this approach one defines three categories of: 1) above a certain threshold (pathogenic evidence), 2) below a different threshold (benign evidence) and 3) in between the two thresholds (no evidence). Here we are using REVEL analysis of the same 22 pathogenic and 27 benign RYR1 variants. The REVEL output is numerical, which facilitates trying different thresholds with a gene of interest and the known variants. After trying several different sets of thresholds we set an upper threshold of >0.85 and a lower threshold of <0.50. To derive Odds ratios for these data we use what is called a 2 x k odds ratio calculator (several are available online – I use https://www.scistat.com/statisticaltests/likelihoodratios.php ) where we set k=3, since the data will be in the form of a 2 x 3 table (note that we have to insert a 1 in the cell for the 'Path;<0.50' cell even though REVEL did not report any of the 22 known path variants with a score below the lower threshold. This is important because using zero in any of the cells makes the calculations problematic. If we put a zero in that cell, the LR for the <0.50 category would come out as zero, which means infinite or perfect classification, which is, as we all know, impossible. By putting a one in this cell of the LR table I am in effect making my data worse, and thus giving a more conservative, but more useful, estimate of the likelihoods).

| | Path | Benign | LR+ | 95% CI |
|---|---|---|---|---|
| ≥0.85 | 19 | 1* | 23.13 | 3.35-159.96 |

* One could further subdivide this into more than three categories using a 2 x 4 or 2 x 5 table. The yet more sophisticated approach would be to perform linear regression on the data and generate LR+ outputs directly from the REVEL readout for each variant and then add that in to the Tavtigian et al equation 5 above, but that is beyond the scope of this analysis.

| | | | | |
|---|---|---|---|---|
| 0.50-0.85 | 3 | 8 | 0.457 | 0.137-1.526 |
| ≤0.50 | 1* | 19 | 0.064 | 0.009-0.443 |

The LR+ for a readout of >0.85 would therefore be 23.13:1, above strong. Note that the 95% confidence interval calculated here ranges from above supporting to above strong. The LR for the <0.50 category is 0.064:1 in favor of pathogenicity, which is 15.63:1 in favor of benign. This is above moerate, although the 95% CI lies between 2.26:1 benign (between supporting and moderate) to 111:1 benign, which is between strong and very strong.

The intermediate range of REVEL output (0.5-0.85) gives an LR of 0.46:1 pathogenic, or 2.19:1 benign. Note here that the 95% CI ranges from 0.137:1 pathogenic (7.30:1 benign) to 1.53:1 pathogenic. This should give one pause that it is not good evidence either way – if the 95% CI broadly flanks 1.0 (no evidence) you are in a middle gray zone and should not use the evidence at all. This makes intuitive sense – there are thresholds for evidence such that the extremes are used as evidence (one end for benign and the other for pathogenic) and the middle range is not.
